# Multiple decisions about one object involve parallel sensory acquisition but time-multiplexed evidence incorporation

**DOI:** 10.1101/2020.10.15.341008

**Authors:** Yul HR Kang, Anne Löffler, Danique Jeurissen, Ariel Zylberberg, Daniel M Wolpert, Michael N Shadlen

## Abstract

The brain is capable of processing several streams of information that bear on different aspects of the same problem. Here we address the problem of making two decisions about one object, by studying difficult perceptual decisions about the color and motion of a dynamic random dot display. We find that the accuracy of one decision is unaffected by the difficulty of the other decision. However, the response times reveal that the two decisions do not form simultaneously. We show that both stimulus dimensions are acquired in parallel for the initial ~0.1 s but are then incorporated serially in time-multiplexed bouts. Thus there is a bottleneck that precludes updating more than one decision at a time, and a buffer that stores samples of evidence while access to the decision is blocked. We suggest that this bottleneck is responsible for the long timescales of many cognitive operations framed as decisions.

## Introduction

Decisions are often informed by several aspects of a problem, each guided by different sources of information. In many instances, these aspects are combined to support a single judgment. For example, an observer might judge the distance of an animal by combining perspective cues, binocular disparity and motion parallax. In other instances, the aspects are distinct dimensions of the same object. For example, the animal’s distance and its identity as potential predator or prey. The former problem of cue combination (***Jacobs, 1999**; **Ernst and Banks, 2002***) is a topic of study in what has been termed the Bayesian vision or the Bayesian Brain (***Knill and Pouget, 2004***). The latter is the subject of this paper. It arises in a wide variety of problems whose solutions depend on identifying a set of conjunctions such as the ingredients of a favorite dish, or when one must make multiple judgments, or decisions, about the same stimulus.

The neuroscience of decision-making has focused largely on perceptual decisions, contrived to promote the integration of noisy evidence over time toward a categorical choice about one stimulus dimension. A well studied example is a decision about the net direction of motion of randomly moving dots. In such binary decisions (e.g., left or right), behavioral and neural studies have shown that humans and monkeys accumulate noisy samples of evidence and commit to a choice when the accumulated evidence reaches a threshold (***Ratcliff, 1978**; **Palmer et al., 2005**; **Gold and Shadlen, 2007**; **Stine et al., 2020***). The framework has been extended to more than two categories (e.g., ***Churchland et al. 2008**; **Bogacz et al. 2007**; **Ditterich 2010***) but it remains focused on a common stream of evidence bearing on a single stimulus feature. Less is known about how multiple streams of evidence are accumulated for a multidimensional decision (***Lorteije et al., 2015***). Given the parallel organization of the sensory systems, one might expect all available evidence to be integrated simultaneously. However, there are also reasons to suspect that two decisions cannot be made in parallel. This is based on a variety of experiments that expose a “psychological refractory period” (PRP; ***Welford 1952***). When participants are asked to make two decisions in a rapid succession, it appears that the second decision is delayed until the first decision is complete (***Pashler, 1994***). Based on such observations, it has been argued that there is a structural bottleneck in the response selection step, such that only one response can be selected at a time (***Sigman and Dehaene, 2005***).

Here we develop a task in which the participant views one visual stimulus and makes two decisions about the same object. The stimulus comprises elements that give rise to two streams of evidence bearing on their motion and color, and the participant must decide on both aspects and report the combined category. The task was designed to allow participants to integrate both streams of evidence simultaneously from the same location in the visual field and to require just one response. We show that, even in this situation, the two streams of evidence are accumulated one at a time. We show that this seriality arises despite the parallel access of the visual system to both streams. We suggest that seriality is explained by a bottleneck between the parallel acquisition of evidence and its incorporation into separate decision processes. We elaborate a model of bounded evidence accumulation, used previously to explain both the speed and accuracy of motion (***Palmer et al., 2005***) and color decisions (***Bakkour et al., 2019***), and show that these accumulations must occur in series. The results have implications for a variety of psychological observations concerning sequential vs. parallel operations, and they address the fundamental question of why mental processes take the time they do.

## Results

We studied variants of a perceptual task that required binary decisions about two properties of a dynamic random dot display. Human participants decided the dominant color and direction of motion in a small patch of dynamic random dots (Fig. 1). The stimulus is similar to one introduced by Mante et al. (***2013***), who studied the problem of gating when making a decisions about only a single dimension, either color or motion. On each video frame, each dot has a probability of being colored blue or yellow and it has another probability of being plotted either at a displacement Δ*x* relative to a dot shown 40 ms earlier or, alternatively, at a random location in the display. We refer to the probability of a displacement as the coherence or strength and use its sign to designate the direction. We use an analogous signed probability for the color coherence or strength (see Methods). Participants reported their answer by making an eye or hand movement to select one of four choice targets. We refer to this as a double-decision and refer to the two aspects as stimulus dimensions. We employed several variants of this basic task in our study.

**Figure 1.**
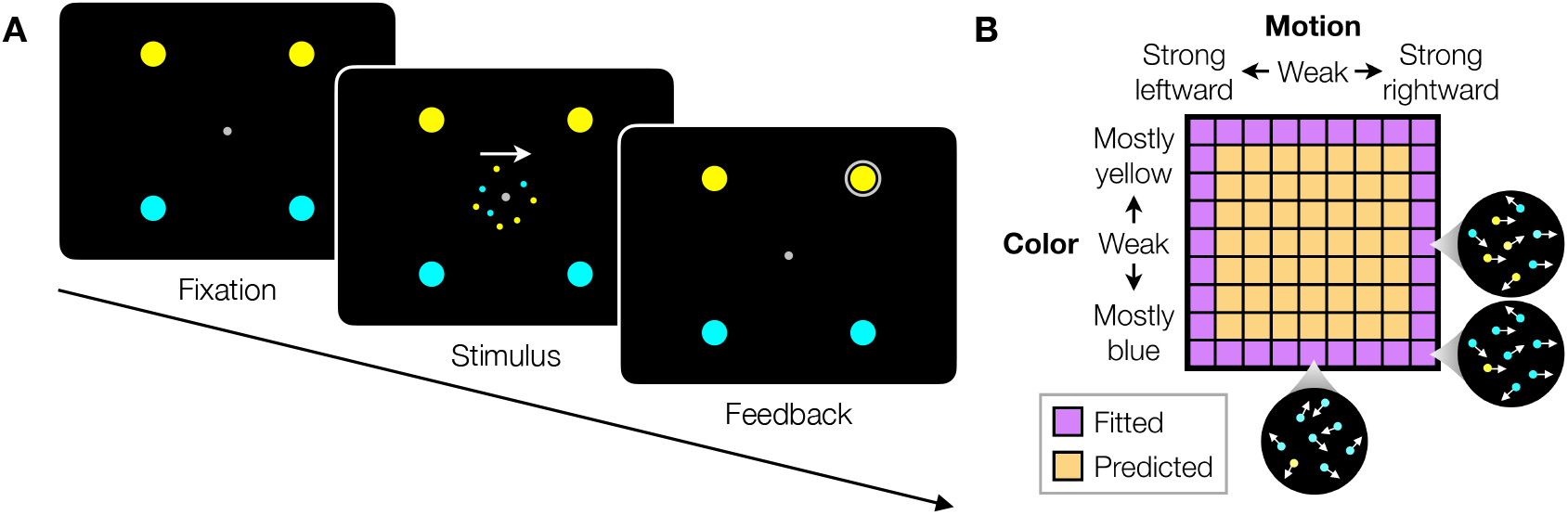
Double decision task. **A**. Timeline of the behavioral task. Participants first fixated a gray dot at the center of the screen. A dynamic random dot stimulus was displayed and the participant was asked to judge the overall motion direction and the dominant color (the arrow is for visualization purposes only and was not presented to the subject). They reported this double decision by selecting one of four targets to indicate motion direction (left and right target for leftward and rightward motion, respectively) and color (top yellow vs. bottom blue targets). The response was deemed correct when both motion and color judgments were correct. Participants received auditory feedback as to whether they were correct and the correct target was also indicated by a white ring. Across the experiments the targets could be indicated with an eye movement or a hand movement, either when the participant was ready to report (reaction time) or when the dot display was extinguished (experimenter-controlled duration). **B**. Motion and color strengths were varied independently across trials, represented by a matrix of combinations of difficulty levels (here shown for the eye reaction time experiment with 81 combinations;see Methods). Insets illustrate typical motion and color for three of the conditions. Correctness was assigned randomly when the coherence was zero. For the combinations shown in purple, at least one stimulus dimension was at its strongest value (easiest). For some analyses, the data from these combinations are used to fit a model, which is evaluated by predicting the data from the remaining combinations (yellow).

A brief précis of the experimental results may be helpful. We first present the main finding using a free response paradigm, what we term *double-decision reaction time*. It demonstrates no interference in choice accuracy—that is, the difficulty of the color decision does not affect the accuracy of motion decisions, and vice versa—but critically, the double decision time is the sum of the two single decision times. The analysis suggests that the motion and color decisions are not formed at the same time. This establishes the prediction that with brief stimulus presentations, successful color decisions ought to be attained at the expense of motion, and vice versa—that is, choice interference. We then test this prediction and fail to confirm it. We show that color and motion can be acquired in parallel but are unable to update the decision simultaneously. This confirms the response selection bottleneck predicted by Pashler (***Fagot and Pashler, 1992***) and it implies the existence of buffers (***Sperling, 1960**; **Kamienkowski and Sigman, 2008***), where sensory information can be held before it updates a decision variable—the accumulated evidence for color or motion.

The combination of a buffer and serial updating leads to a revised prediction that interference in accuracy should occur over a narrow range of stimulus viewing duration, controlled by the experimenter. We confirm this prediction, showing that there is no interference at short viewing times, but that there is a narrow regime of the stimulus duration in which accuracy on one dimension suffers because a limited amount of deliberation time needs to be shared with the other dimension, which reconciles conflicting observations of parallel and serial patterns of decision-making in the literature (e.g., ***Schumacher et al. 2001**; **Tombu and Jolicœur 2004***). We then introduce a bimanual version of the task which affords direct report of both the color and motion termination times. It confirms the assumption that the double-decision time is the sum of two sequential sampling processes, each with its own stopping time, and it shows that the color and motion decisions compete before the first decision terminates. This implies some form of time-multiplexed alternation. In the last experiment we ask participants to judge whether the motion in a pair of patches are the same or different and find that this binary decision also exhibits additive decision times. Finally, we introduce a conceptual model of the double-decision process that serves as a platform to connect the computational elements with known and unknown neural mechanisms.

### Double-decision reaction time

Participants were asked to judge both the net direction (left or right) and dominant color (yellow and blue) of a patch of dynamic random dots and to indicate both decisions with a single movement to one of four choice targets (Fig. 1A). Different groups of participants performed the task by indicating their choices with an eye movement or a reach (see Fig. 5A). On each trial the strength and direction of motion as well as the strength and sign of color dominance were chosen independently, leading to 81 (9×9 eye) or 121 (11×11 arm) combinations. The single movement furnished two decisions and one reaction time (RT). Participants were given feedback that the decision was correct if the motion and color were both correct (see Methods).

Fig. 2A & B shows choices and mean RT as a function of stimulus strength for the eye and hand tasks, respectively. The graphs in the left column of each panel show the data plotted as a function of motion strength and direction. Each color on this graph corresponds to a different difficulty of the other dimension (i.e., color). Similarly, the graphs in the right columns show the data plotted as a function of color strength and dominance; the uninformative dimension, motion, is shown by color. Unsurprisingly, the proportion of rightward choices increased as a function of the sign and strength of the motion coherence, and the proportion of blue choices increased as a function of the sign and strength of color coherence. The slopes of these logistic functions supply an estimate of sensitivity. The striking feature of these graphs is that sensitivity to variation in the stimulus along each dimension is unaffected by the difficulty along the uninformative dimension. This is evident from the superposition of the colored data points. It is also supported by a logistic regression analysis, which favored a choice model in which the sensitivity along one dimension is not influenced by the stimulus strength along the other dimension (ΔBIC = 23 and 22 for motion and color in the eye task, respectively; ΔBIC = 37 and 50 for the hand task; positive values are support for the regression model of Eq. 12 without the *β*_3_ term). It implies that the two stimulus dimensions do not interfere with each other. This is consistent with the well established idea that color and motion are processed by parallel, independent channels (***Carney et al., 1987***). However, another possibility is that the two dimensions do not interfere because they are not processed simultaneously but serially.

**Figure 2.**
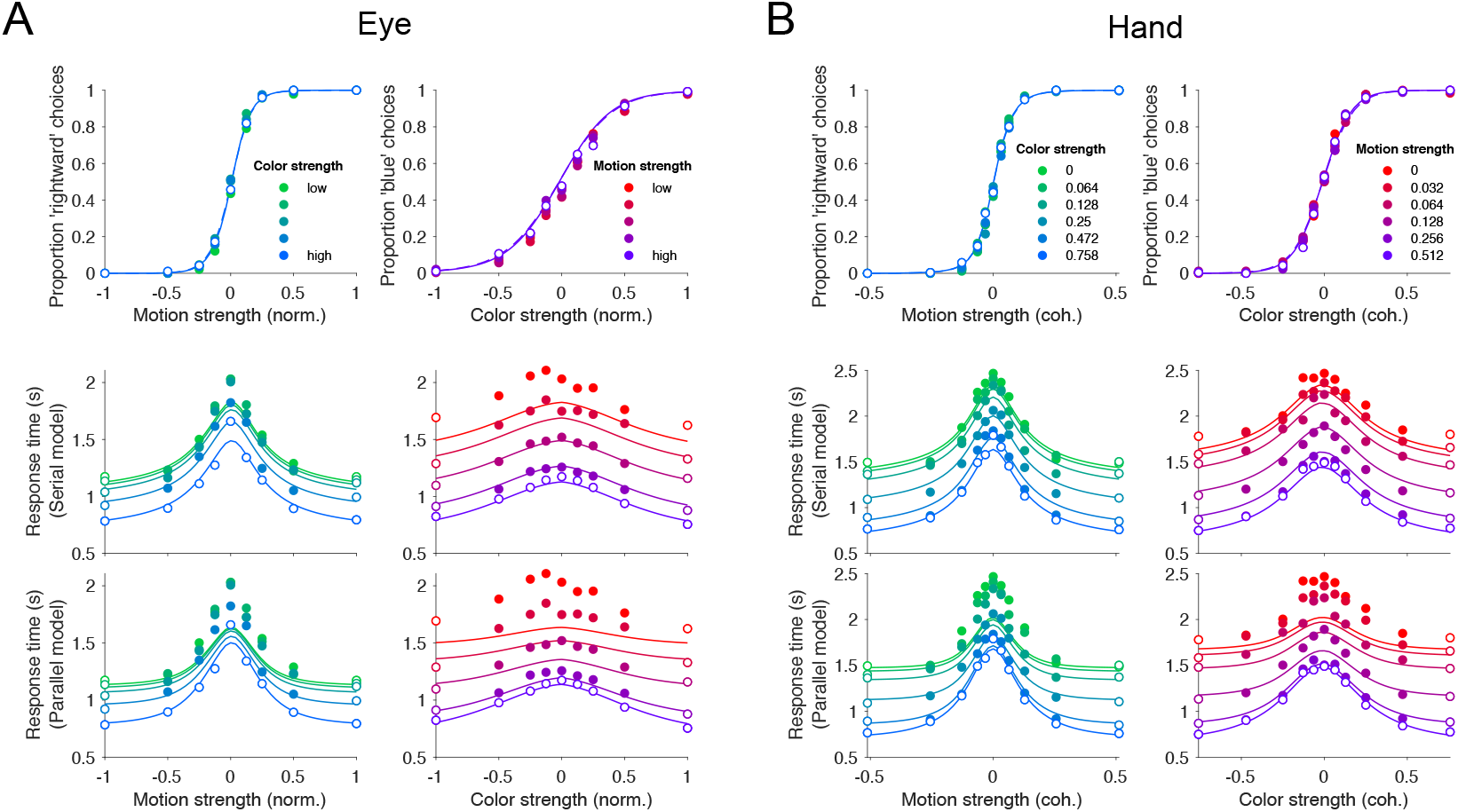
Double-decisions exhibit additive response times but no interference in accuracy. Participants judged the dominant color and direction of dynamic random dots and indicated the double-decision by an eye movement (A) or reach (B) to one of four choice-targets. All graphs show the behavioral measure (proportion of choices, top row; mean reaction time, rows 2 and 3) as a function of either signed motion or color strength. Positive and negative color strength indicate blue- or yellow-dominance, respectively. Positive and negative motion strength indicate rightward or leftward, respectively. Colors of symbols and traces indicate the difficulty (unsigned strength) of the other stimulus dimension (e.g., color, for the graphs with abscissae labeled “Motion strength”). Symbols are combined data from three participants (Eye) and eight participants (Hand). Open symbols identify the conditions used to fit the parallel and serial models. These are the conditions in which at least one of the two stimulus strengths was at its maximum (purple shading, Fig. 1B). The models comprise two bounded drift-diffusion processes, which explain the choice and decision times as a function of either color or motion. They differ only in the way they combine the decision times to explain the double-decision RT. For the serial model, the double decision time is the sum of the color and motion decision times. For the parallel model the double-decision time is the longer of the color and motion decisions (see Methods). Smooth curves are the predictions based on the fits to the open symbols. Both models predict no interaction on choice (top row). The serial predictions (middle row) are superior to the parallel model. Data are the same in the lower two rows. Stimulus strengths in A were not identical for the 3 participants and were combined to a common scale. **Figure 2–Figure supplement 1.** Statistical comparison of the drift diffusion model under serial vs. parallel rules. **Figure 2–Figure supplement 2.** Comparison of parallel and serial rules applied to reaction time distributions. **Figure 2–Figure supplement 3.** Statistical comparison of parallel and serial rules applied to reaction time distributions. **Figure 2–Figure supplement 4.** Mean reaction time for parallel and serial rules applied to the reaction time distribution analysis. **Figure 2–Figure supplement 5.** Mean reaction time for parallel and serial rules applied to reaction time distribution analysis with the fit-prediction approach

Indeed, the RTs support this serial hypothesis. The reaction times, plotted as a function of either motion or color, exhibit inverted U-shapes, such that longer reaction times are associated with the most difficult stimulus strength and the fastest with the easiest. In contrast to the choice functions, the uninformative dimension—that is, with respect to the dimension of the abscissa—affects the scale of these RTs, giving rise to a stacked family of inverted U-shaped functions. The more difficult the other dimension, the longer the RT.

We attempted to explain the choice-RT data in Fig. 2 with models of bounded evidence integration (e.g., drift-diffusion; ***Ratcliff 1978**; **Palmer et al. 2005***). Such models provide excellent accounts of choice and RT on the motion-only and color-only versions of these tasks (***Palmer et al., 2005**; **Bakkour et al., 2019***). To explain the double-decision data set we pursued two variants of these models under the assumption that motion and color are processed in parallel or in series. The curves in Fig. 2 are a mixture of fits and predictions. To fit the data (open symbols), we used all trials in which at least one of the dimensions was at its strongest level (32 purple conditions in Fig. 1B for the eye task and 40 conditions for the hand task). We used these fits to predict the data from the remaining conditions (49 amber conditions for the eye, Fig. 1B and 81 for the hand; filled symbols, Fig. 2). Both models are consistent with no interference in the choice functions. Thus the fit to the 32 or 40 conditions supplies all the predicted choice functions.

The models can be distinguished on the basis of the RT data. For an experiment with only a single dimension (e.g. motion), the RT is the sum of the amount of time that evidence is integrated to reach a terminating bound (the decision time, *T*_m_ or *T*_c_, for motion and color choice respectively) plus additional time for sensory and motor delays, termed the non-decision time (*T*_nd_). If the color and motion decisions are made in parallel, then the total decision time should be determined by the slower process (max[*T*_m_,*T*_c_]), whereas if the decisions are made serially, the total decision time would be determined by the sum of the two decision times (*T*_m_ + *T*_c_). In both cases, we expect both motion and color strengths to affect the RT. In the serial case, an increase in the difficulty of color, say, should augment the total RT by the same amount for all motion strengths, giving rise to stacked functions of the same shapes (solid curves, middle row, Fig. 2A,B). In the parallel case, an increase in the difficulty of color should augment the total RT by an amount that depends on the difficulty of motion (solid curves, bottom row Fig. 2A,B). The color dimension is likely to determine the total RT when motion is strong, but it has less control when the motion is weak. The logic should produce stacked bell-shaped functions that pinch together in the middle of the graph. The data are better explained by the serial predictions (e.g, large mismatches when both dimensions are weak). Formal model comparison provides strong support for the serial models overall (geometric mean of Bayes factor across participant and task combinations: > 10^39^) and for 9 out 11 participants individually (***Figure 2–Figure Supplement 1***).

We pursued a second approach to compare serial and parallel integration strategies, focusing specifically on the decision times. Unlike the fits to choice-RT, this method uses each participant’s choices as ground truth. It considers only the distribution of RTs and attempts to account for them under serial and parallel logic. Instead of diffusion models, we estimated the marginal distributions for each 1D decision time and the four *T*_nd_ distributions (for each choice) with gamma distributions. For the serial case the predicted RT distributions are established by convolution of the marginal single-dimension distributions and the distribution of *T*_nd_. For the parallel case the marginals are combined using the max logic, and the result is convolved with the appropriate distribution of *T*_nd_ (see Methods). ***Figure 2–Figure Supplement 2*** shows fits to the reaction time distribution for the more informative conditions for the serial and parallel models. The model comparisons, based on all the data, yield “decisive” support (***Kass and Raftery, 1995***) for the serial processing of motion and color (geometric mean of Bayes factor for participant and task combinations > 10^18^ with all participants’ individually supporting the serial rule; ***Figure 2–Figure Supplement 3***). We also display the mean RTs derived from the fits in the same format as Fig. 2 (***Figure 2–Figure Supplement 4*** & ***Figure 2–Figure Supplement 5***).

The finding favors additive decision times, from two independent decision processes, each with its own termination rule. However, it does not discern the nature of the serial processing (e.g., whether they alternate or one is prioritized). We will consider this issue later.

### Brief stimulus presentation

The results from the double-decision RT experiment support sequential updating of two decision variables, which represent accumulated evidence for the motion and color choices. If this is true, it leads to a straightforward prediction. If the stimulus duration is not controlled by the decision maker but by the experimenter, and if it is brief, then the two stimulus dimensions would compete for the limited processing time, and we ought to observe choice-interference. We therefore conducted a second experiment in which we limited the duration of the stimulus viewing time to just 120 ms. We know from previous experiments with 1D tasks that performance increases with stimulus durations greater than one half second (***Kiani et al., 2008**; **Waskom and Kiani, 2018***). Thus it is reasonable to assume that performance accuracy would suffer if it is not possible to make use of the full 120 ms of evidence for both motion and color. We predicted that sensitivity to both color and motion should be worse on the double-decision task than on color-only and motion-only versions of the identical task.

To our surprise, double-decisions were just as accurate as their 1D controls (Fig. 3A). We also observed no change in the sensitivity to color across the range of motion difficulties, and vice versa (ΔBIC = 9 and 10 for motion and color choices, respectively, in support of no interaction; Eq. 11, *H*_0_: *β*_3_ = 0). This suggests that evidence for color and motion were acquired simultaneously, in parallel, and without interference. Further support for this conclusion is adduced from an analysis of the stimulus information used to make the decisions—what is known as psychophysical reverse correlation or kernel (***Beard and Ahumada, 1998; Okazawa et al., 2018***). Fig. 3B displays the degree to which trial-by-trial variation in the noisy displays influences the choice (see Methods). It shows that these stimulus fluctuations influenced choices almost identically in the double-decision task and 1D controls.

**Figure 3.**
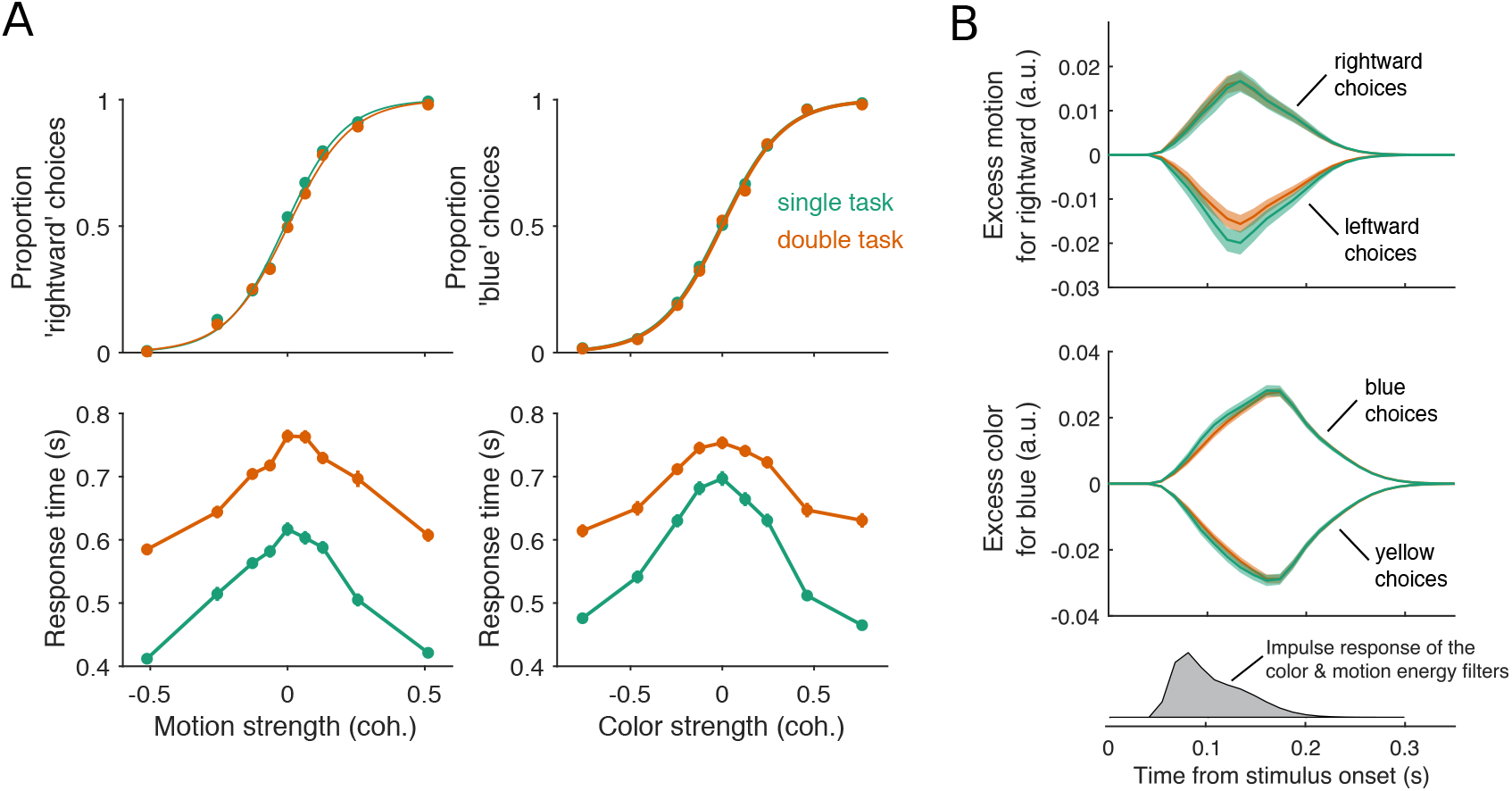
Parallel acquisition and serial incorporation of a brief color-motion pulse. Participants completed a short-duration variant of the double-decision task in which the stimulus was presented for only 120 ms. They also performed blocks in which they were asked to report only the color or only the motion direction (single decision in which they could ignore the irrelevant dimension). Data from double- and single-decision blocks are indicated by color. **A**. Choice probability and response times for single and double decision blocks. *Top-left*, proportion of rightward choices as a function of motion strength. Top-right, proportion of blue choices as a function of color strength. The solid lines are logistic fits. They are nearly identical for single- and double-decisions. *Bottom row*, Response times for the single- and double-decisions plotted as a function of motion strength (left) and color strength (right). For the double decisions, these are the same data plotted as a function of either the motion or color dimension. (Three participants performed a total 9,959 double-decision and 12,527 single-decision trials). Data points show the average response time as a function of motion or color coherence, after grouping trials across participants and all strengths of the “other” dimension (i.e., color, *left*; motion, *right*). Error bars indicate s.e.m. across trials. Although the stimulus was presented for only 120 ms, response times were modulated by decision difficulty. Importantly, response times were longer in the double-decision task than in the single-decision task. **B**. Psychophysical reverse correlation analysis. *Top*, Time course of the average motion information favoring rightward, extracted from the random-dot display on each trial, that gave rise to a left or right choice. Shading indicates s.e.m. *Middle*, Time course of the average color information favoring blue, extracted from the random-dot display on each trial, that gave rise to a blue or yellow choice. The shaded area indicates the s.e.m. across trials. The similarity of the green and orange curves indicates that participants were able to extract the same amount of information from the stimulus when making single- and double-decisions. *Bottom*, Impulse response of the filters used to extract the motion and color signals (see Methods). They explain the long time course of the traces for the 120 ms duration pulse.

At first glance, the observation seems to be at odds with our interpretation of the double-decision RT experiment, which provided strong support for serial processing, primarily in the pattern of RTs. Here, the entire stimulus stream lasts only 120 ms, which is less than a typical saccadic latency to a bright spot. Nevertheless, participants exhibited variation in the time of their responses as a function of stimulus strength (Fig. 3A, bottom panels) and these response times were surprising long. The fastest were ~300 ms longer than the stimulus (*RT* > 400 ms). Importantly, they are approximately 100-200 ms longer in the double-decisions than in single decisions. It is difficult to make too much of this observation, because the participants might have procrastinated for reasons unrelated to the dynamics of the decision process. However, procrastination would not explain the difference between the two conditions. As parallel acquisition of the 120 ms color and motion take the same amount of time as acquisition of either of the streams alone (by definition), the extra time in the double decision is probably explained by serial incorporation of evidence into the two decisions. This observation also implies the existence of buffers that store the information from one stream as it awaits incorporation into the decision.

Our results so far suggest that color and motion information are acquired in parallel but are incorporated into the decision in series. We therefore wondered if the same schema might apply to the double-decision RT task. For this to hold, some kind of alternation must occur such that segments of one or the other stimulus stream is not incorporated into its decision. Suppose, for example, that at t=120 ms, motion information had been incorporated into decision variable *V*_m_, and color information had been stored in a buffer. Suppose further that motion continues to update the decision variable, *V*_m_, until it reaches a termination bound at *t = T*_m_, and only then can the buffered color information be incorporated into decision variable, *V*_c_. From then on color information could update *V*_c_ until this decision terminates. In this imagined scenario, the color information between 0.12 < *t* < *T*_m_ is not incorporated in the decision.

One might also imagine two alternatives to the latter part of this scenario. In both, the information from color continues to update the buffer (but not *V*_c_) throughout the motion decision without loss. Then at *t = T*_m_ either (*i*) all the information about color is incorporated immediately into *V*_c_ or (*ii*) the buffered information is incorporated in *V*_c_ over time (e.g., as if the recorded color information is played back). The first alternative is equivalent to the parallel model that is inconsistent with the data. The second scenario, implausible as it may seem, implies the color decision is blind to the color information in the display during the playback of the recorded color information (i.e., *T*_m_ < 2*T*_m_). These alternatives are not intended as serious models but to convey two general intuitions. First, if there is a buffer at play in the 2D reaction time task then it must take time for the buffered information to be incorporated, or the RTs would have conformed to the parallel logic. Second, if the duration of the buffer is finite, when both 1D processes require more processing time than the duration of the buffer, there will be portions of the color and/or motion stimulus that do not affect the decision.

One might therefore ask why the second point does not lead to a reduction in sensitivity (or accuracy) in color, say, when motion is weak and competes with color for processing time. The answer is that when the decision maker controls the termination of the decision, they can compensate the missing information by collecting more until the level reaches the same terminating bound. This leads to a straightforward prediction. If the experimenter controls the termination of the evidence stream, then missing portions of the color and/or motion stimulus might impair performance, especially when the other stimulus dimension is weak.

### Variable stimulus duration

We therefore predicted that under conditions in which the experimenter controls the viewing duration, there is an intermediate range of viewing durations, greater than 120 ms and less than the average RT of difficult double decisions, where we might observe interference in sensitivity. To appreciate this prediction, it is essential to recognize that when the experimenter controls viewing duration of a random dot display, the decision maker applies a termination criterion, as they do in free response (RT) experiments (***Kiani et al., 2008***). There is no overt manifestation of this termination, although it can be identified by introducing perturbations to the stimulus (see also ***Kang et al. 2017***). Before such termination, accuracy improves by the square root of the stimulus viewing duration 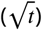 as expected for perfect integration of signal-plus-noise. In a double-decision, when the two decision processes are splitting the time equally, the accuracy of each should only improve by 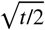. However, when one process terminates, the rate of improvement of the other process should recover, until that process reaches its terminating bound. The model predicts a range of stimulus strengths and viewing durations in which interference in accuracy ought to be evident. It also predicts that the range and degree of interference might depend on which stimulus dimension the participant prioritizes. Here we set out to test this prediction.

Two participants performed this variable stimulus duration task in 12-16 sessions. The task was identical in structure to the brief-duration experiment. However, stimuli were presented at fixed durations ranging from 120 to 1200 ms (in steps of 120 ms). Only three levels of difficulty were used for each dimension: one easy and two difficult coherence levels (adjusted individually to yield 80% and 65% accuracy, respectively; see Methods). All 6 × 6 combinations of motion × color coherences were presented.

Fig. 4 shows the sensitivity to motion and color as a function of stimulus duration, when the other stimulus dimension was easy or difficult. The sensitivity is the slope of a logistic fit of the motion (or color) choices to the three levels of difficulty (see Methods). Notice that for both participants, there is no difference in the slopes at the shortest stimulus duration (120 ms), consistent with the findings above. However both participants exhibited lower sensitivity at intermediate durations when color choices were coupled with difficult motion. This difference implies an interference. It is less compelling, if present at all, when motion choices are coupled with difficult color. This pattern in which motion difficulty affects color sensitivity but not vice-versa is consistent with participants prioritizing one decision over the other. This would arise if participants consistently monitored the motion stream first and turned to color after the motion decision terminated. In this case the difficulty of the color would not affect the decisions for motion, but harder motion would take longer to terminate thereby leaving less time for color processing. We therefore used a model in which one decision was prioritized over another by including a parameter that determined the probability that motion would be processed first. We also included a parameter that controls the duration of the stimulus streams that can be held in the buffer. This is, effectively, the amount of stimulus information that can be acquired in parallel. The best fits of the model, shown by the smooth curves (Fig. 4A) suggest the buffer capacity of 40-200 ms worth of stimulus information (Fig. 4B & ***Figure 4–Figure Supplement 1***) and prioritization of motion on approximately 80-96% of trials. Had the buffer capacity been tiny, the model would be purely serial and if the buffer duration was very large, then the model would be parallel. Both such buffer capacities provide very poor fits to the data (***Figure 4–Figure Supplement 2***).

**Figure 4.**
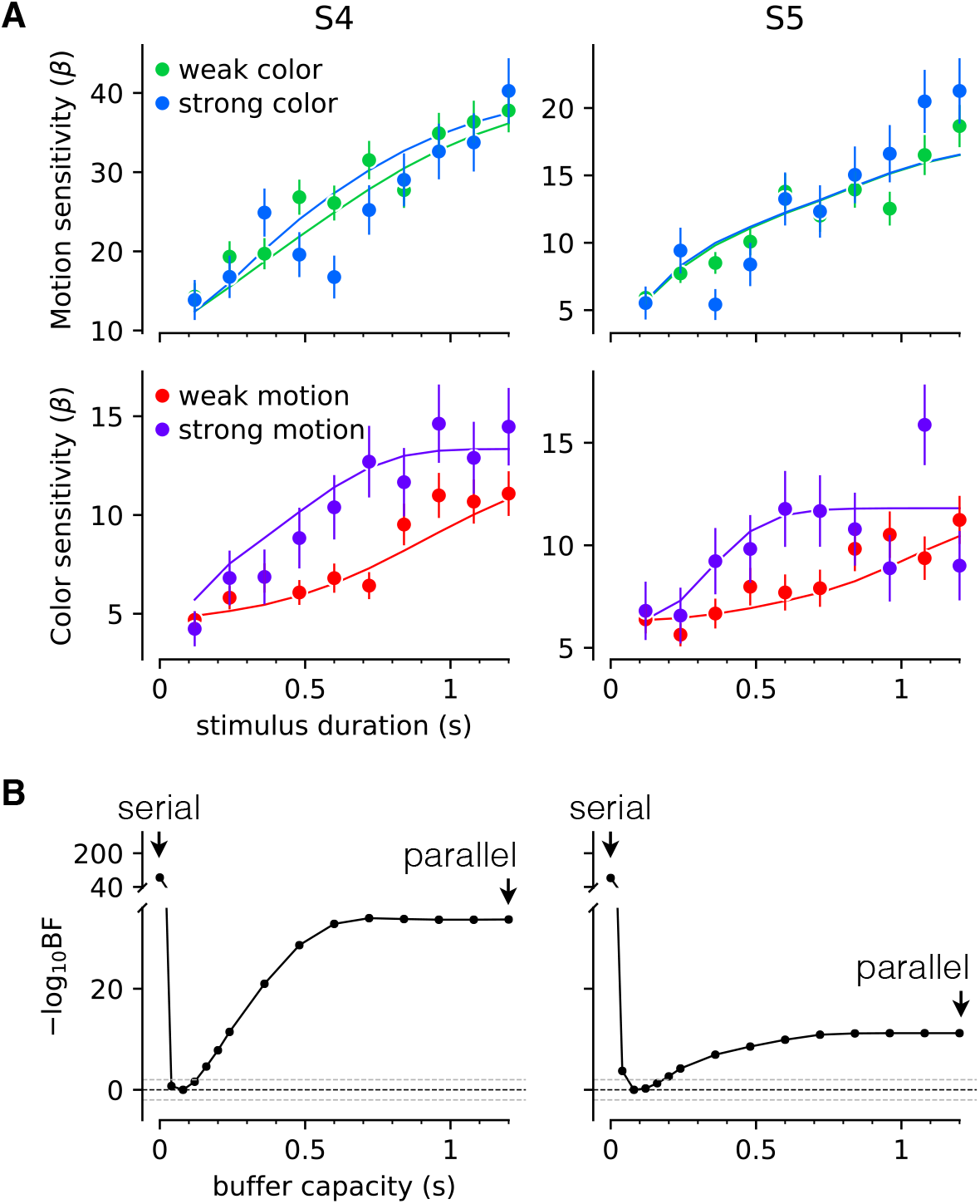
Interference in choice accuracy can be elicited at intermediate viewing durations. Two participants (columns) performed the color-motion double-decision task with a random dot display that varied in duration between 120 and 1200 ms. **A**. *Top*, Motion sensitivity as a function of stimulus duration and color strength. Symbols are the slope of a logistic fit of the proportion of rightward choices as a function of signed motion strength, for each stimulus duration. Data are split by whether the color strength was strong (blue) or weak (green). Error bars are s.e. *Bottom*, Analogous color-sensitivity split by whether the motion strength was strong (purple) or weak (red). Curves are fits to the data from each participant using two bounded drift diffusion models that operate serially after an initial stage of parallel acquisition, here termed the buffer capacity. During the serial phase, one of the dimensions is prioritized until it terminates. The prioritization favored motion for both participants (*p*_motion-1st_ = 0.80 and 0.96, for participants S4 and S5, respectively). **B**. Negative log likelihood of the model fits as a function of the buffer capacity, relative to the model fit at 80 ms capacity. The model is equivalent to a purely serial model, when the buffer capacity is zero, and to a purely parallel model when the buffer capacity exceeds the maximum stimulus duration. Negative log likelihoods were computed for a discrete set of buffer capacities (black points). Black dashed lines are at Bayes factor = 1 (log_10_BF = 0). Gray dashed lines show where the Bayes factor = 100 (“decisive” evidence for the best fit model compared to the models above the line; ***Kass and Raftery 1995***). **Figure 4–Figure supplement 1.** Parameter recovery analysis **Figure 4–Figure supplement 2.** Fits to the choice data with strictly serial and parallel models

The findings therefore support our prediction and in doing so, they support the hypothesis that a common principle explains the double decisions ranging from a tenth to at least two seconds and whether this duration is controlled by the experimenter or by the decision maker. Namely, there is parallel acquisition but serial incorporation of color and motion into the double-decision process. The interference in choice accuracy demonstrated in this experiment is the only example of choice interference in our study. It is remarkably elusive, because it can be observed only for stimulus durations for which three conditions are satisfied: (*i*) the duration of the stimulus is long enough that parallel acquisition is no longer possible; (*ii*) the duration of the stimulus is short enough that accuracy on one dimension would benefit from additional sensory evidence; (*iii*) the duration should support termination of the other dimension for strong but not weak stimuli. The interference is also deceptive. It is explained by a competition for processing time, not by an interaction affecting the fidelity of the sensory streams themselves. It is an example of resource sharing (***Tombu and Jolicœur, 2002**, **2005**; **Kahneman, 1973***), but the resource is time, specifically.

### Separate effectors (bimanual)

There are two important features of the serial model: the existence of two decision variables that are terminated independently, and that these accumulations are not updated at the same time but in series. A limitation in the experiments so far is that we had access to the completion of the double decision but not to the completion of each component. Therefore, we could only speculate about which decision completed first and when. Without knowledge of the first decision time, we cannot tell how often a participant switched between updating the motion and color decision variables. For example, the prioritization considered in the previous section could arise by completing one decision before deliberating on the second or by alternating back and forth on a schedule that allocates more time to motion. Therefore, we conducted an experiment in which participants indicated their choice and RT for each stimulus dimension using separate effectors.

The eight participants who performed the unimanual version of the double-decision RT task also performed a bimanual version of the same task (Fig. 5A). In the unimanual version, participants used a handle to move a cursor to one of four targets that simultaneously communicated color and motion decisions. In the bimanual version, participants indicated their motion decision by moving one of the handles in a left/right direction and indicated their color decision with a forward/backward movement of the other handle. Participants were encouraged to independently indicate their color and motion decisions. To facilitate this, they received extensive training, consisting of blocks in which one of the stimulus dimensions was set at its easiest level. Both the order of the tasks (unimanual and bimanual) and the hand assignments (left/right × color/motion) were balanced between the participants (see Methods).

**Figure 5.**
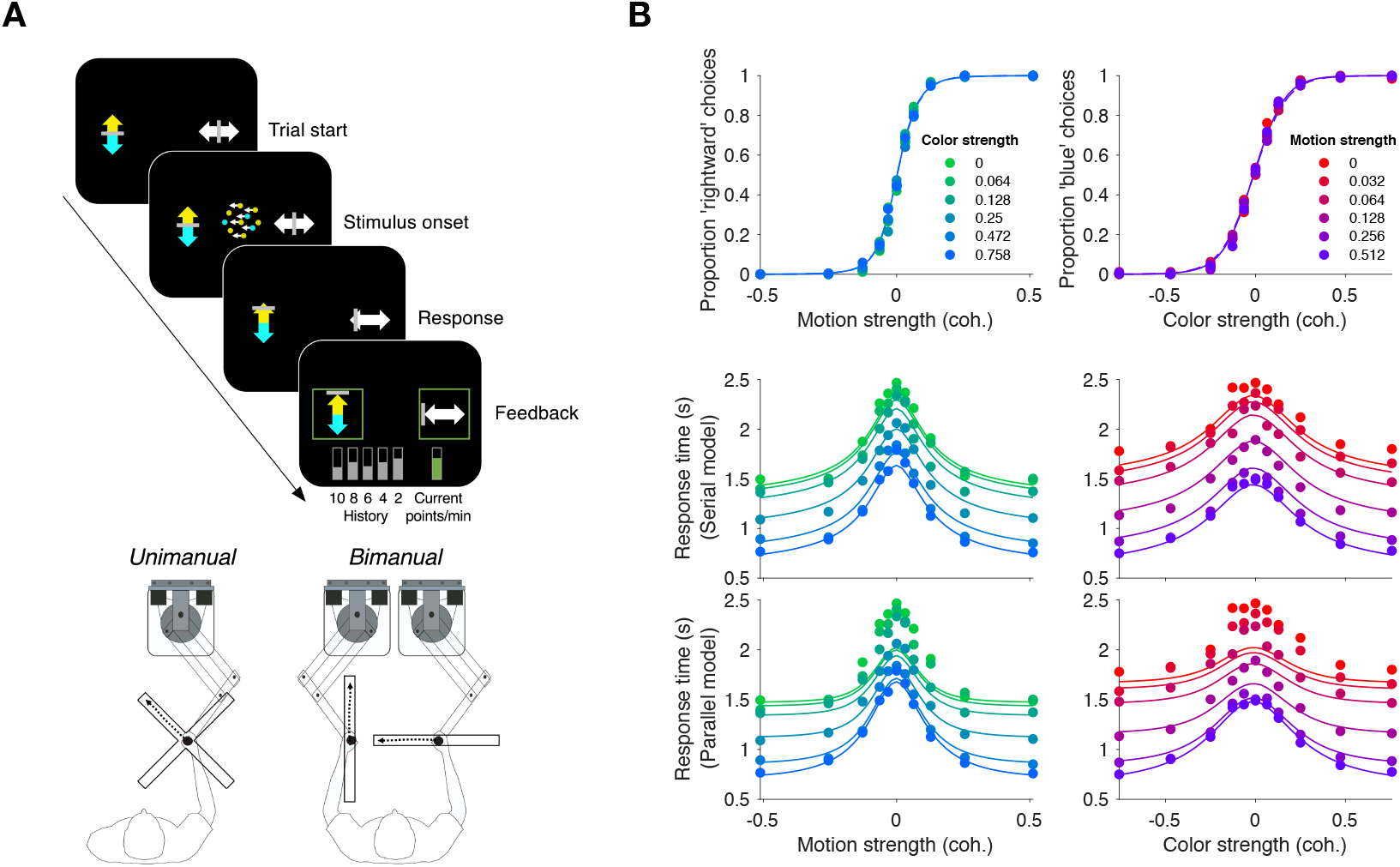
Replication of double-decision choice-reaction time when the decisions are reported with two effectors. **A**. Participants performed the color-motion double-decision choice-reaction task, but indicated the double-decision with either a unimanual movement to one of four choice-targets or a bimanual movement in which each hand reports one of the stimulus dimensions (N=8 participants performed both tasks in a counterbalanced order). In both conditions the hand or hands were constrained by a robotic interface to move only in directions relevant for choice (rectangular channels). The display was the same across unimanual and bimanual tasks with up-down movement reflecting color choice and left-right movement reflecting motion choice. A scrolling display of proportion correct was used to encourage accuracy. In the unimanual trials both choices were indicated simultaneously. However, in the bimanual trials each choice could be indicated separately and the dot display disappeared only when the second hand left the home position. **B**. Choice proportions and double-decision mean RT on the bimanual task. The double-decision RT on the bimanual task is the latter of the two hand movements. The data are plotted as a function of either signed motion or color strength (abscissae), with the other dimension shown by color (same conventions as in Fig. 2). Solid traces are identical to the ones shown in Fig. 2 for the unimanual task, generated by the method of fitting the conditions containing at least one stimulus condition at its maximum strength and predicting the rest of the data. They establish predictions for the bimanual data from the same participants. The agreement supports the conclusion that the participants used the same strategy to solve the bimanual and unimanual versions of the task. **Figure 5–Figure supplement 1.** Choice and double-decision reaction time for the bimanual responses **Figure 5–Figure supplement 2.** Model-free comparison of performance in the unimanual (blue) vs. bimanual (red) task.

Before tackling the questions that motivate the bimanual experiment, we first ascertained whether participants used the same strategy to make bimanual double-decisions as they did on the unimanual version. It seemed conceivable that by using separate hands to indicate the motion and color decisions, participants could achieve parallel decision formation, for example, as a pianist reads the treble and bass staves with the left and right hands, typically. We therefore conducted a model comparison similar to that of Fig. 2. To fit the models, we used the color and motion choice on each trial along with the second response time (D_2nd_) regardless of whether it was to indicate direction or color. This allows us to fit models that are identical to those used in the unimanual task (Fig. 2). In the bimanual task, the final RTs (RT_2nd_) are well described by the fits to the unimanual double-decision RTs (Fig. 5). We illustrate this in two ways. In the figure, the solid traces are not fits to the bimanual data; they are fits to the unimanual data shown in Fig. 2B. Clearly the choice probabilities and response times displayed in the bimanual task are well captured by the model fit to the unimanual task. The actual fits are shown in ***Figure 5–Figure Supplement 1***, and model comparison favors the serial over the parallel model for seven of the eight participants (***Figure 2–Figure Supplement 1***). Importantly, the participants’ behavior was strikingly similar in the unimanual and bimanual versions of the task.

The similarity between the two versions of the task is also supported with a model-free analysis. In ***Figure 5–Figure Supplement 2*** we superimpose the accuracy and the reaction times for the unimanual and bimanual tasks. There is an almost perfect overlap between these two aspects of choice behavior, providing further support for a common set of processes operating in both versions of the task. It provides direct evidence for two termination events, as assumed in our model fits. This rules out a class of models of the double decision as a race among four accumulations for each of the color-motion combinations, what we term targetwise integration, as these models posit only one double-decision time.

The bimanual task allows us to distinguish between two variants of the serial model that were not distinguishable in the unimanual task. In the first variant, the *single-switch* model, the decision maker only switches from one decision to the next when the first decision is completed. Thus the decision that terminates first (D_1st_) is the one that is evaluated first, and only then the other decision is evaluated. In the second variant, the *multi-switch* model, the decision maker can alternate between decisions even before finalizing one of them. If little time is wasted when switching, these two models make similar predictions for the response time in the unimanual task: the response time will be the sum of the two decision times plus the non-decision latencies. However, the models make qualitatively different predictions for how the response time for D_1st_ depends on the difficulty of the other decision.

The single-switch model predicts the response time for D_1st_ is independent of the difficulty of the decision reported second (D_2nd_). That is because D_2nd_ is not evaluated until the first decision is completed. The prediction of the multi-switch model is less straightforward. Suppose that in a given trial the motion decision is easy and the color decision is difficult. If the color was reported first, the motion was probably not evaluated at all before committing to D_1st_, since if it had been evaluated it would most likely have ended before the color decision. In contrast, if both dimensions were difficult, which decision was reported first is largely uninformative about the number of alternations between color and motion that occurred before committing to the first decision; since both decisions take longer to complete, it is possible that both have been evaluated before one of them terminated. Therefore, the multi-switch model predicts that the first decision takes longer the more difficult the other decision is: when D_2nd_ is easy, it is more likely that it was not considered before committing to the D_1st_ decision and thus the average response time is shorter.

To disambiguate between the single-switch and multi-switch models, we fit both models to the data from the bimanual task. First, we fit a serial model identical to that of Fig. 2 to the data from the bimanual task. We used the same procedure as in Fig. 2; that is, we ignore RT_1st_ and fit RT_2nd_ and the choices given to the two decisions. Then, we used three additional parameters to attempt to explain RT_1st_. These parameters are the average time between switches (*τ*_Δ_), the probability of starting the trial evaluating the motion decision (*p*_motion-1st_), and the non-decision time for the first decision 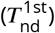. These parameters only affect RT_1st_; they do not influence RT_2nd_ nor the choices made for the two decisions. The three parameters were fit to minimize the mean-squared error between the models’ predictions and the data points (Fig. 6; Table 3). The single switch model is a special case of the multi-switch model where *τ*_Δ_ is very large (i.e., longer than the slowest first decision time).

**Figure 6.**
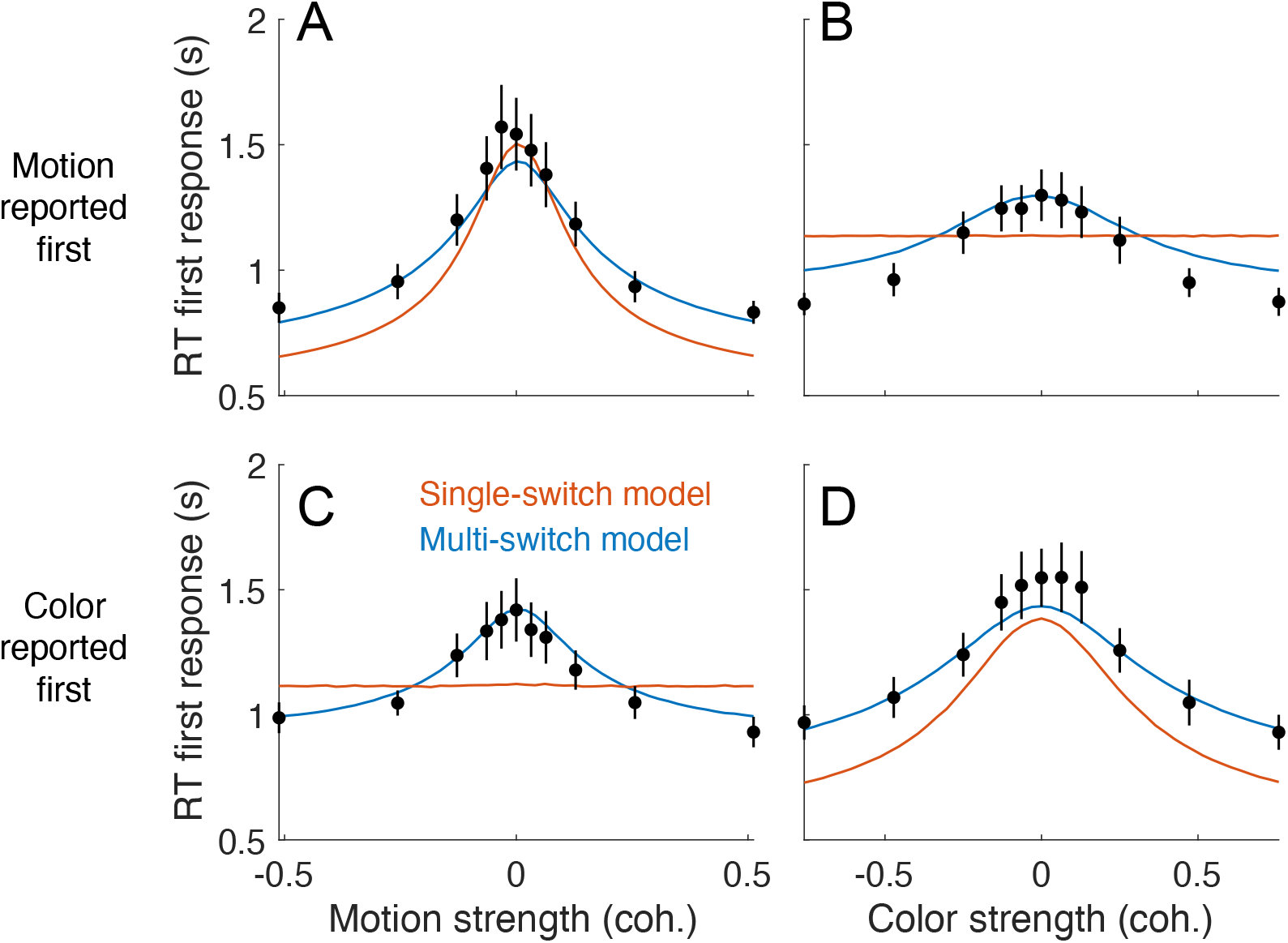
First response times in the bimanual task suggest multiple switches in decision updating. For bimanual double decisions, participants indicate two RTs per trial. Whereas up to now we have only considered the RT corresponding to completion of both color and motion decisions, the analyses in this figure concern the RT of the first of the two. Symbols are means ± s.e. (N=8 participants). Curves are fits to single- and multi-switch model (colors). **A**. RT as a function of motion strength when motion was reported first. **B**. RT as a function of color strength when motion was reported first. **C**. RT as a function of motion strength when color was reported first. **D**. RT as a function of color strength when color was reported first. In panels A and D, the 1st response corresponds to the stimulus dimension represented on the abscissa. The data exhibit the expected pattern fast RT when the stimulus is strong and slow RT when the stimulus is weak (i.e., near 0). This would occur if the serial processing of motion and color ensued one after the other (single-switch) or with more than one alternation (multi-switch), although the latter provides a better account of the data. In panels B and C, the 1st response corresponds to the stimulus dimension that is not represented on the abscissa. Here the single-switch model fails to account for the data. If there were only one switch and color terminates first, then the strength of motion is irrelevant, because all processing time was devoted to color. Similarly, if there were only one switch and motion terminates first, then the strength of color is irrelevant, because all processing time was devoted to motion.

The model comparison provides clear support for multiple switches. Fig. 6 shows the average response time for the decision reported first (RT_1st_), split by whether the first decision was color or motion, and grouped by either color or motion strength. Both the single- and multi-switch models provide a good explanation of the RT_1st_ when grouped as a function of the coherence of the decision that was reported first (Fig. 6, panels A and D). However, only the multi-switch model could explain the interaction between RT_1st_ and the coherence of D_2nd_ (Fig. 6B and C). The data shows that RT_1st_ is longer when D_2nd_ is more difficult, and this effect was well explained by the multi-switch model. Unlike what is seen in the data, the single-switch model predicts that RT_1st_ should not vary with the coherence of D_2nd_ (as depicted by flat lines in panels B and D). Because we fit the models for each participant individually, we can analyze the frequency of alterations predicted by the model with multiple switches. For one of the participants, the best-fitting interswitch interval was higher than the slowest decision time, and thus the model was no different from the single-switch model. For the other 7 participants, alternations were sparse: the average inter-switch interval was 920±290 ms (mean ± s.e.m. across participants).

To summarize, the bimanual version of the double-decision task allowed us to infer not only that the two dimensions were addressed serially, but that people may alternate between both attributes of the stimulus in a time-multiplexed manner. The model suggests that alternations were sparse, as if the participants considered one decision for a few hundred milliseconds, and switched temporarily to the other decision if they found no conclusive evidence about the first.

### Double decision with binary response

Upto nowwe have observed serial decision making when participants had to provide two answers— that is, four possible responses. A possible concern is that the reason we observed the serial pattern of double-decisions was that it required a quaternary response. We therefore designed a task that involves a double decision but only a binary choice. Two participants were asked to report whether the net direction in two patches of random dots were the same or different (Fig. 7A). The two motion stimuli were presented to the left and right of a central fixation cross Fig. 7A. The direction (up or down) and strength of motion were controlled independently in the two stimuli.

**Figure 7.**
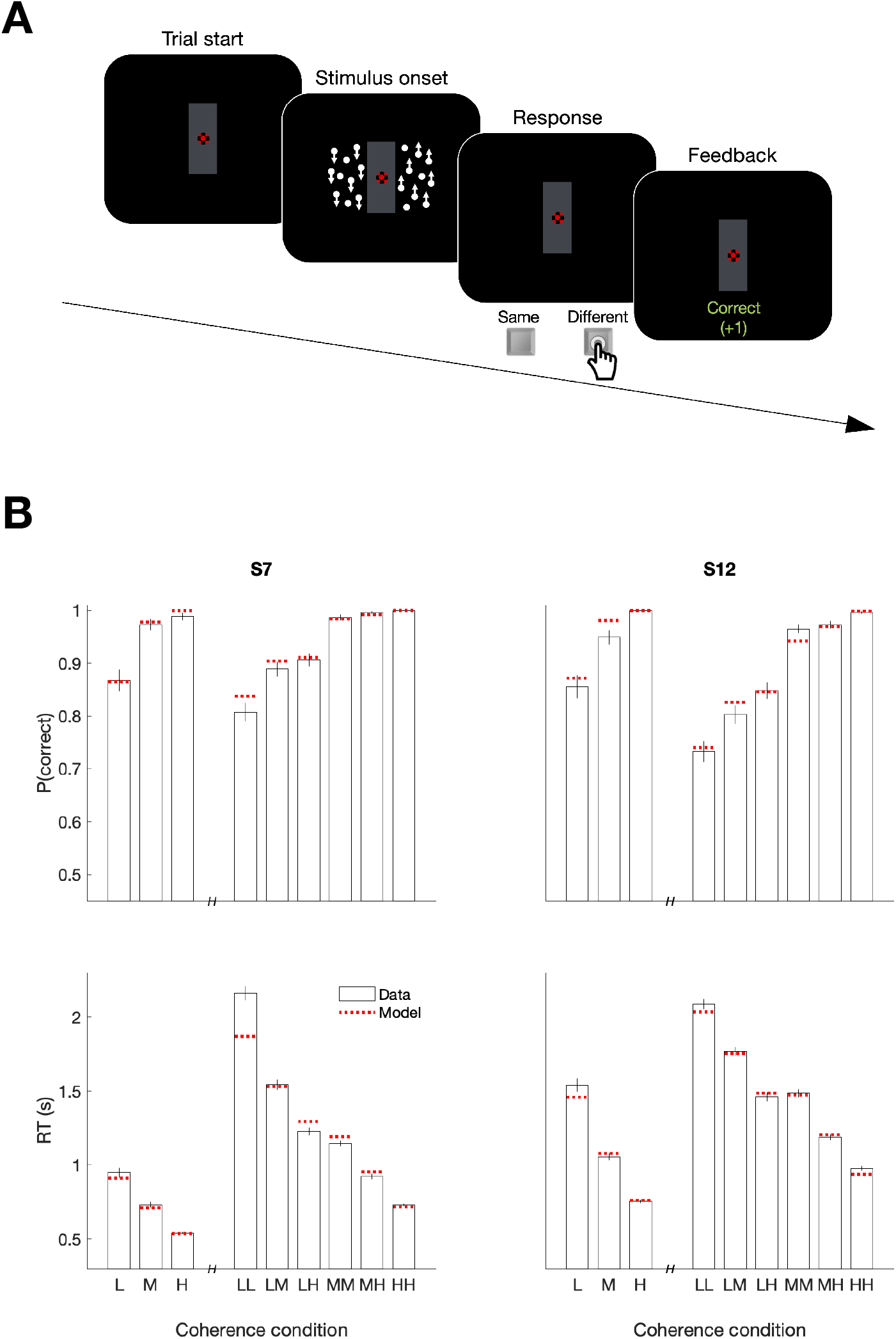
Serial decision making in a Same vs. Different task. **A**. Task. Two dynamic random dot motion displays were presented in rectangular patches to the left and one to the right of a central fixation cross. The direction and motion strength were randomized from trial to trial and between the patches (up or down × three motion strengths). Participants judged whether the dominant direction of the left and right patches is the same or different and indicated the decision when ready by pressing a response key with their left or right index finger. At the end of each trial, participants received feedback. In a separate block, participants also performed a 1D direction discrimination task in which only one patch of random dots was displayed. **B**. *Top*, Proportion of correct choices as a function of the level of absolute motion strength (L = low;M = medium; H = High). *Bottom*, Reaction times for each level of motion strength. The first three bars represent the direction task where only a single motion stimulus was presented. The six bars on the right of each plot represent the same-different task. Horizontal red lines are fits of a serial drift-diffusion models to the means. Only correct trials were included for RT analyses. **Figure 7–Figure supplement 1.** Comparison of parallel and serial rules applied to reaction time distributions in the Same vs. Different task

Both participants exhibited accuracy-RT functions that depended on the difficulty of both motion stimuli. Fig. 7B shows proportion of correct choices plotted as a function of the coherences for both the 1D (up-down) and 2D (same-difference) trials. The RTs associated with same-different judgment were almost twice as long as the RTs from a 1D direction judgment. Part of this difference might be attributed to the conversion from two direction judgments to the same-different response, but that should not depend on difficulty and it is hard to reconcile this with the magnitude of the difference. Instead they suggest additive decision times. The horizontal red lines in Fig. 7B are fits to a drift diffusion model that assume the 2D same/different decision is formed from two 1D direction decisions. We constrained the fits to share the same sensitivity to motion strength (see Methods, Eq. 2).

To compare serial and parallel accounts of these extended reaction times, we used the same strategy as in ***Figure 2–Figure Supplement 2*** which attempts to account for the observed RT distributions as combinations of underlying 1D decision times and the non-decision time. This analysis provides strong support for the serial account (***Figure 7–Figure Supplement 1C***; BF > 10^7^ for both participants). Like the color-motion task, there is every reason to assume that the acquisition of evidence from the two patches of random dots occurs in parallel. Yet once again, the pattern of RTs supports serial incorporation into the double decision. The use of a binary response in the same-different task also rules out the possibility that the long decision time in our 2D experiments are explained by the doubling of alternatives (Hick’s law; ***Hick 1952**; **Luce 1986**; **Usher et al. 2002***).

### Parallel acquisition with serial incorporation model

Taken together, the results from our five experiments suggest that the prolongation of RTs in double decisions is the result of serial integration of evidence during the decision-making process, independent of the modality of choice implementation and number of response options. Parallel acquisition of the two sensory streams followed by serial incorporation into decision variables reconciles the findings of the short duration experiment with those of the double-decision RT experiment. The variable duration and bimanual experiments suggest that (i) parallel acquisition and serial incorporation is not limited to the short duration experiment and (ii) serial alternation of color and motion can occur before one process terminates. Here we attempt to incorporate these features into a common framework intended to illuminate how this might work in terms that relate to the neurobiology of perceptual decision making. We will proceed by illustrating the steps that underlie the acquisition of evidence samples, their temporary storage in buffers, and their incorporation into the decision variables that govern choice and the two decision times. We first make the case for the buffer using a simulated trial from the short duration experiment. We then elaborate the diagram to account for the serial pattern of decision times when the stimulus duration is longer.

Consider the example in Fig. 8A of a process leading to a decision in the short duration task. Suppose that visual processing of the 120 ms motion stream gives rise to a single sample of evidence that captures the information from the brief pulse, and the same is true for the color stream. These samples of evidence are acquired in parallel and placed in buffers, where they can be stored temporarily. The values in these buffers may be thought of as latent instructions to a cortical circuit to update a decision variable (*V*_m_ or *V*_c_) by some amount (Δ*V*_m_ and Δ*V*_c_). While the samples can be acquired simultaneously, only one sample can update the corresponding decision variable at a time. This is the bottleneck. One of the samples must be held (buffered) until the other update operation has cleared. If motion is the first to be updated, then *V*_c_ cannot be updated until the circuit receiving the motion-update instruction has received it (green arrow). This takes some amount of time, *τ*_ins_ (for **ins**truct). The update instruction is realized by an integrator with a time constant (*τ*_v_ = 40 ms) leading to slow cortical dynamics (red and blue traces).

**Figure 8.**
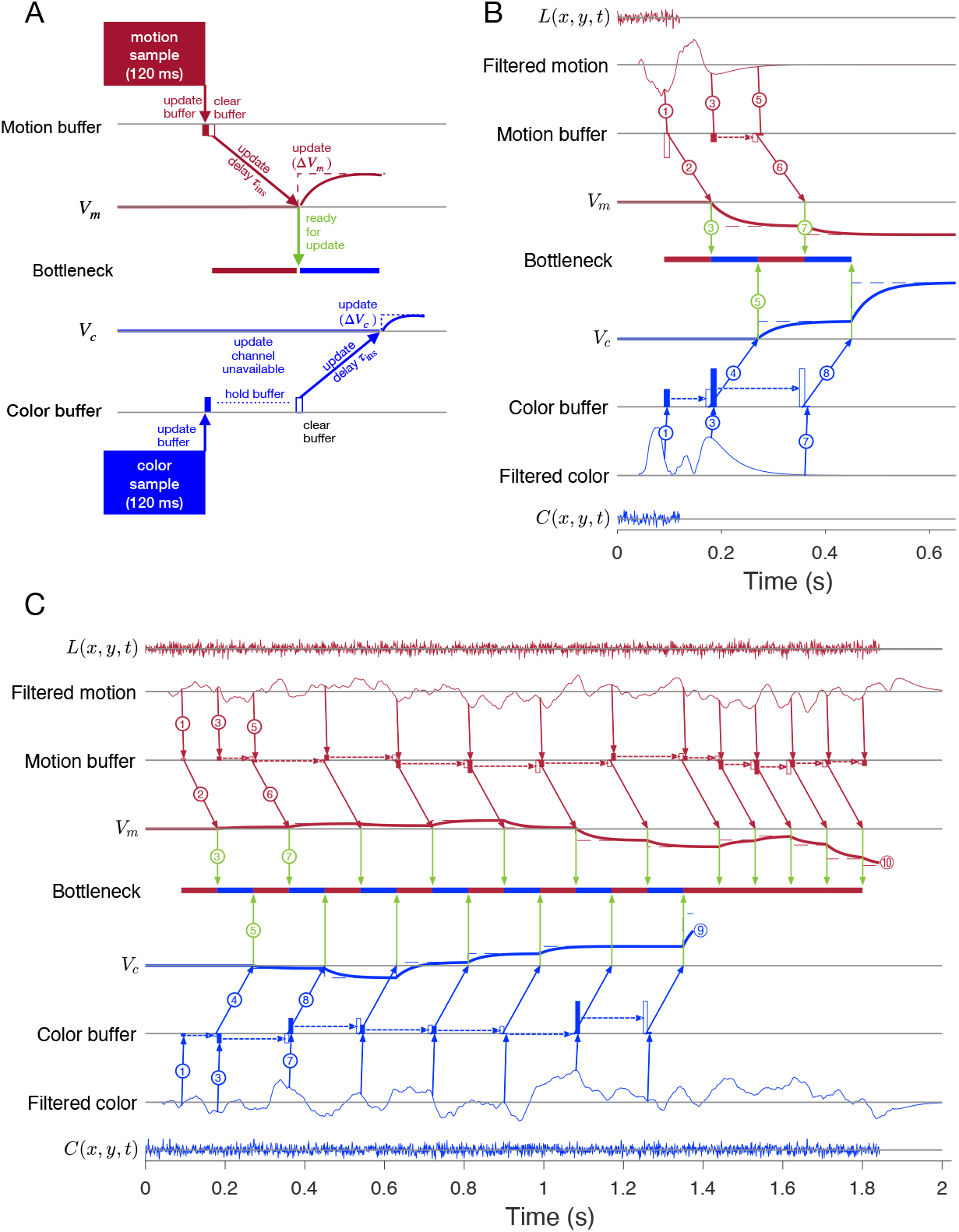
Parallel acquisition of evidence and serial updating of two decision variables. An elaborated drift diffusion model permits reconciliation of the serial processing implied by the double-decision choice-RT experiment and the failure to observe interference in choice accuracy when the color-motion stimulus is restricted to a brief pulse. The main components of the model are introduced in panel A and elaborated in panels B and C. In all panels, red and blue indicate motion and color processes, respectively. **A**. Simulated trial from the short duration experiment. Information flows from top to bottom graphs for motion; and from bottom to top graphs for color. Time is left to right. The evidence from both color and motion is extracted from the 120 ms random dot stimulus in parallel. Both can be stored temporarily in separate buffers (filled rectangles), which send an instruction to the circuits representing the respective decision variables in their persistent firing rates. The instruction is to change the firing rate by an amount (Δ*V*_m_ or Δ*V*_c_). This latency from clearance of the sample from the buffer to receipt of the Δ*V* instruction takes time (*τ*_ins_, black diagonal arrows), and this is followed by the realization of the instruction in the evolving firing rates of cortical neurons (smooth colored curves). In the example, the *V*_m_ is the first to update. A central bottleneck precludes updating *V*_c_. The bottleneck is cleared when the Δ*V*_m_ instruction is received by the circuit that represents the motion decision variable (green arrow). This allows the buffered evidence for color to update *V*_c_. Open rectangle represents clearance of the buffer content, which occurs immediately for motion and after a delay for color in this example. Dashed lines associated with decision stage show the instructed change in the decision variable (Δ*V*_m_ and Δ*V*_c_). Smooth colored curves show the evolution of the decision variables. **B**. Elaboration of the example in panel-A. The boxes representing the 120 ms stimulus are replaced by the two outer rows: (i) raw luminance and color data stream, £(x,y,r) and C(x,y,r), respectively, represented as biased Wiener processes (duration 120 ms); (⊓) filtered evidence streams containing the relevant motion (right minus left) and color (blue minus yellow) signals. The filters introduce a delay and smoothing. The filtered signals can be sampled by the buffer every *τ*_s_ ms, so long as the buffer is available (i.e., empty). The bottleneck shows the process that is accessing the update channel. Other than the first sample, the prioritization is equal and alternating. Only one process can update at a time. Circled numbers identify the key events described in Results. Events sharing the same number are approximately coincidental. **C**. Example of a double-decision in the choice-reaction time task. The first eight steps parallel the logic of the process shown in panel B. The decision variables then continue to update serially, in alternation, until *V*_c_ reaches a terminating bound ((9). The decisions then continues as a 1D motion process until *V*_m_ reaches a terminating bound (¢§). Bound height is indicated by (9 and Note that the sampling rate is the same as it was in the parallel phase, whereas during alternation it was half this rate for each dimension.

In this example, each buffer receives all the information available in the stimulus. Were there additional samples in the stimulus, the motion buffer would be ready to receive another sample when it sends its content, whereas the color buffer cannot be updated until it is cleared, *τ*_ins_ later. The bottleneck is between the buffer and the update of the decision variable, more specifically, the initiation of the dynamic process that implements this update in a cortical circuit. In this case there is no consequence beyond a delay, because there is no more evidence from the stimulus after 120 ms.

Fig. 8B elaborates the diagram in panel A using another trial from the short duration experiment. We now represent the transformation of sensory data to evidentiary samples by applying a stage of signal processing to the raw luminance and color data, *L*(*x, y, t*) and *C*(*x, y, t*). These functions are just shorthand for the noisy spatiotemporal displays. The motion filter is meant to capture the impulse response of direction selective simple and complex cells in the visual cortex (***Movshon et al., 1978b**,a; **Adelson and Bergen, 1985**; **Britten et al., 1993**; **DeAngelis et al., 1993***), and we assume a similar operation on the stimulus color stream. They are also shorthand for a difference signal, such as right minus left and blue minus yellow. The filtering introduces a delay and a smearing of these streams. While the motion filters must sample the *L*(*x,y,t*) at rates sufficient to support the extraction of fast fluctuations and fine spatial displacement, the neurons ultimately pool these signals nonlinearly over space and time (***Britten et al., 1993**; **Zylberberg et al., 2016***). These are the signals represented by the maroon filter traces in the Fig. 8B. This is the convolution of *L*(*x, y, t*) and the function in Fig. 3B (bottom). The same filter is applied to *C*(*x, y, t*) to make the filtered color traces (blue). Importantly, for purposes of integrating the information in the color-motion random dot displays, 11 Hz sampling (*τ*_s_ = 90 ms) is sufficient. Notice that the filtered representation lasts longer than the stimulus. Therefore, in this case the decision is based on at least two samples of evidence per sensory stream.

The buffers acquire their first samples at *τ*_s_ = 90 ms (Fig. 8B, arrows 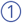 & 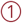) The motion buffer is cleared as soon as it is acquired (arrow 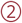) and thus begins to impact *V*_m_ 90 ms later (i.e., *τ*_ins_).

We set *τ*_ins_ = 90 ms mainly to simplify the figure (but see below). Thus it is only at *t* = 180 ms (i.e., *τ*_s_ + *η*_ns_) that the instruction arrives to update *V*_m_. This unblocks the bottleneck 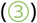, thereby allowing the first color sample to be cleared from its buffer (open rectangle, 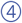). This permits acquisition of a second color sample (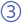 and filled blue rectangle, *t* = 180 ms). Notice that the second motion sample is also acquired at *t* = 180 ms, that is, *τ*_s_ after the first acquisition (and its immediate clearance). The first color sample instructs *V*_c_ *τ*_ins_ after it was cleared (*t* = 270 ms), which unblocks the bottleneck (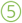). Because we are assuming alternation in this example, this leads to the second update of *V*_m_ 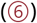. With the motion buffer available, it would be possible to obtain a third sample from the motion stream at *t* = 270 ms, but the filtered signal has decayed to zero, and we assume extinction of the stimulus is registered by the brain by this time to terminate sampling. Upon receipt of Δ*V*_m_, the bottleneck is lifted (*t* = 360 ms; 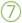) and the second color sample is cleared from its buffer (*t* = 360 ms) to instruct *V*_c_ 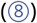. There is no signal left to integrate, and the decision is made based on the signs of *V*_m_ and *V*_c_. Thus the decision is based on simultaneous (parallel) acquisition of two samples of evidence, which are incorporated serially into their respective decision variables.

The exercise helps us appreciate how a stream of evidence lasting only 120 ms could lead to a double-decision 400-600 ms later (Fig. 3A). It also illustrates the compatibility of parallel acquisition and serial incorporation into the decisions, and it suggests that serial processing is imposed at the step between buffered samples and incorporation into the decision variables. This is the “response selection” bottleneck hypothesized by Harold Pashler (1994) and others (e.g., ***Marti et al. 2012***; see Discussion).

The idea extends naturally to double-decisions that are extended in time. Fig. 8C illustrates a simulated double-decision in a free response task. The double-decision is made once both decision variables reach their terminating bounds. The example follows the same initial steps as the short duration experiment, except that when the 2^nd^ motion and color samples are cleared from their respective buffers, they are replaced with a 3^rd^ sample. Notice that beginning with the third motion sample, the interval to the next sample has doubled (180 ms), because the example posits regular alternation (for purposes of illustration only; see below). This longer interval begins with the 2^nd^ sample. From that point forward, until the color decision terminates, the streams are effectively undersampled. Decision processes ignore approximately half of the evidence supplied by the stimulus. This is because both streams supply independent samples of evidence at a rate greater than 5.5 Hz (i.e., an interval of 180 ms).

In the example, it is *V*_c_ that reaches the bound first (*T*_c_ ≈ 1.4 s; 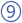). There may be no overt behavior associated with this terminating event, as in the eye and unimanual reaching tasks, but direct evidence for this termination is adduced from the bimanual reaching task.

From this point forward the processing is devoted solely to motion until it terminates at a negative value of *V*_m_ 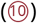. Notice that when the bottleneck clears, there is always a buffered sample ready to be cleared, and this occurs at intervals of *τ*_ins_ = *τ*_s_ =90 ms. The process is now as efficient as a single decision process. Indeed, a simple 1D decision about motion (or color) is likely to involve the same instruction delays and bottleneck. If *τ*_s_ = *τ*_ins_, then like the first sample of motion, all subsequent samples of motion could pass immediately from the buffer to update *V*_m_ without loss of information. The model is thus a variant of standard symmetrically bounded random walk or drift-diffusion (***Laming, 1968**; **Link, 1975**; **Ratcliff, 1978**; **Shadlen et al., 2006**; **Ratcliff and Rouder, 1998**; **Palmer et al., 2005***). It is compatible with the long time it takes for visual evidence to impact the representation of the decision variable in cortical areas like the FEF and LIP (e.g., ~180 ms).

The diagrams in Fig. 8 are intended for didactic purposes, to lay out the need fora buffer and the seriality imposed by a bottleneck between the buffer and the update of the DV in circuits associated with working memory. The values for the delays and time constants (*τ*_x_ terms) were chosen mainly to simplify an already complex diagram, and the same holds for the assumption of strict alternation. The logic does not change if the serial processing were to involve many updates of color or motion before switching to the other dimension. The important assumption is that it takes time to update a decision variable, and during this update there is a bottleneck that precludes another update. Importantly, whether alternating, as in Fig. 8C, or starting one process after completing the other, there is a period of time in which information in the sensory stream is not affecting one of the decisions. This loss is apparent in the additivity of decision times, but it leads to no interference in accuracy in the RT task, because the termination criterion has not changed, and this (and the stimulus strength) determines accuracy. This is the insight that led to the prediction that under certain conditions in which the experimenter controls the duration of the color-motion display, there ought to be interference between color and motion sensitivity (Fig. 4).

## Discussion

In one sense the present study extends the framework of bounded evidence accumulation to more complex decisions composed of the conjunction of two decisions about two distinct features. In another more important sense, the findings highlight a bottleneck in information processing that touches on the very speed of thought. The experimental findings demonstrate that a double-decision about the dominant color and direction of motion of a patch of random dots is formed serially. This is surprising, because color and motion are canonical examples of parallel visual pathways from the retina through the visual and extrastriate visual association cortex, and there are compelling demonstrations of this parallel processing on conscious perception (***Carney et al., 1987**; **Cavanagh et al., 1984**, **1985***). Indeed we confirmed that the color and motion information in the random dot stimulus used here was acquired in parallel. The stimulus was designed to minimize interference or competition for spatial attention. It was restricted to a small aperture in the center of the visual field, and the same individual dots supply the motion and color information. It seems fair to say that the deck was stacked in favor of parallel processing.

Indeed with one notable exception, there was not a hint of an interaction between color or motion on choice performance in our experiments. That is, changing the difficulty of one dimension, say color, did not affect the perceptual accuracy—or more precisely, sensitivity—to the other dimension, say motion. This held over a wide range of difficulties spanning chance to perfect performance. The one exception was when we controlled viewing duration (Fig. 4) and this turns out to be explained by a competition of the two streams for processing time, not by an interaction affecting the fidelity of the sensory streams themselves. Had we attended solely to the choice data, we would have likely concluded that the motion and color decisions were formed in parallel, consistent with 40 years of vision science (***Livingstone and Hubel, 1988**; **Ramachandran and Gregory, 1978***).

Evidence for seriality of the decision process is adduced mainly from the pattern of double-decision reaction times. The RT is the time from the onset of the color-motion stimulus to the initiation of the movement used to indicate the decision: the sum of the time it takes to complete the double decision, plus time delays that are not affected by task difficulty, termed the non-decision time (*T*_nd_). If the color and motion decisions are made in parallel, then the double-decision time is the larger of the two decision times, max[*T*_m_,*T*_c_]. If the decisions are made serially, the double-decision time is the sum, *T*_m_ + *T*_c_. We focused on *max* vs. *sum* distinction using a combination of fitting and prediction. The simplest approach relies only on fits of the double-decision RT distributions (***Figure 2–Figure Supplement 2***) derived from a smaller set of latent 1D decision-time distributions under the appropriate operations for parallel and serial combination (Eq. 7 and convolution, respectively). We found this method the most robustly identifiable—that is, it almost always favors the appropriate generating model—and it is sufficiently powerful to apply to the smaller data sets from individual participants. It reveals “decisive” support (***Kass and Raftery, 1995***) for seriality in all but one of the 11 participants (***Figure 2–Figure Supplement 3***). A drawback of the approach is that it does not constrain the relationship between choice accuracy and decision time. For this we used a variety of bounded drift-diffusion models. These are the fits shown in Fig. 2. Here too, we attempted to contrast the *max* and *sum* logic by predicting the RT distribution for the majority of conditions. We fit the choice-RT data from the subset of conditions in which at least one of the stimulus dimensions was at its strongest level. The fits, under the *max* or *sum* rule, supply the marginal 1D distributions used to predict the RT of the remaining conditions, through application of the same rule. This approach also provides decisive support for the serial model (see ***Figure 2–Figure Supplement 1***).

The strong support for serial processing does not specify where in the processing chain the seriality arises. The answer to this question resolves the apparent contradiction with vision science, and highlights a connection with a body of literature from psychology that addresses the topic of dual task interference, more specifically the psychological refractory period. The key is the short and variable duration experiments (Figs. 3 and 4). If seriality were imposed at the level of sensory acquisition, then when both color and motion are difficult, accuracy on one dimension should come at the expense of accuracy on the other, on average. We did not observe this at short durations, and not for lack of power, as made clear by the interference that was detected at intermediate durations. Nor did we observe any reduction in accuracy compared to single decisions, and there was no difference in the magnitude and time course over which momentary fluctuations of color and motion predicted the individual choices on single- and double-decisions (Fig. 3B). These observation also rule out the possibility that there was interference but it was balanced across trials—that is, a mixture of trials in which successful motion processing impaired color processing on half the trials and successful color processing impaired motion processing on the other half.

If color and motion information are incorporated into the decision serially in the short duration experiment, then there must be a mechanism to store the evidence from at least one of the processes while the other is incorporated into the decision. We refer to this temporary storage as a buffer. There are several reasons to believe that incorporation is serial. First, the response times were longer in the double-decision task than in the single decision task, but the extra time was not associated with improved accuracy on either dimension. Second, the finding was replicated in the variable duration task, which revealed interference of motion on color at intermediate durations, consistent with a serial account.

Thus the short duration experiment demonstrates parallel processing and the necessity of at least one buffer. The results in the variable duration experiment might lead us to entertain the possibility that only color is buffered, because motion was prioritized. However, the bimanual task demonstrates that motion is not always processed first, and both color and motion are processed before the first process terminates. We therefore conclude that there are two buffers which are capable of holding a sample of evidence about color or motion, respectively, while the other dimension is incorporated into the decision. This places the bottleneck between the buffered evidence and the representation of the decision variable. We believe the bottleneck arises because of an anatomical constraint. It is simply impossible to connect in parallel every possible source of evidence with the neural circuits responsible for representing a proposition or plan. As ***Zylberberg et al. (2010)*** theorized, the brain’s routing problem holds the key to why many mental operations operate serially. We will return to this idea after interpreting our results in the context of the neurobiology of decision making. We do this by pursuing the neural correlates of a computational model that supports parallel acquisition of sensory evidence and its serial incorporation into two decisions.

The double decision is formed by two decision processes representing the accumulation of samples of evidence bearing on the dominant color or the dominant direction of motion. Their only interaction is through a competition for access to evidentiary samples, which cannot be supplied to both decision processes at the same time, hence the bottleneck. If one process terminates the other carries on from that time with unfettered access to its momentary evidence. For visual perceptual decisions, parallel acquisition is identified with central visual pathways in the primary and extrastriate cortex. The representation of decision variables is identified with parietal and prefrontal cortical areas and with neurons that exhibit long time scales to support the representation of working memory, planning, and the integration of positive and negative inputs as a function of time. The operations depicted in Fig. 8 are intended to reconcile what is known about the neurobiology of simple 1D decisions with the constraints introduced by the double-decision task. The mathematical instantiation of the model requires only minor modifications of two bounded drift-diffusion processes with temporal multiplexing (see Methods). However the architecture implied by Fig. 8B & C facilitate interpretation of the experimental findings in relation to neural processing.

In the mathematical depiction of drift-diffusion, the momentary evidence is a biased Wiener process. However, in reality the stimulus is not a Wiener process, nor is the representation of momentary evidence by neurons (***Zylberberg et al., 2016***), which arise through application of a transfer function that effectively spreads the impact of a pair of displaced dots over 100-150 ms (Fig. 3B, bottom; ***Adelson and Bergen (1985)***). Thus the neural representation of the motion can be approximated by a process of leaky integration (***Cain et al., 2013**; **Barlow and Tripathy, 1997***). Such smoothing would not be warranted for the detection of fast changes, but it is adequate for a signal that is to be integrated over time. We know less about the filtration of a color difference, but the same logic applies. The conceptual transition from Wiener processes to discrete samples allows us to appreciate the similarity between the accumulation of evidence from movie-like stimuli and the broader class of decisions based on discrete samples of evidence from the environment and memory. This informs hypotheses about the neurobiology, because the sample of evidence ultimately bears on a decision in units of belief or relative value. That is obvious when considering a choice between items on a menu, but it has been camouflaged to some extent in the perceptual decision-making literature. This is in part because the time-integral of a difference in firing rates from right- and left-preferring neurons is the number of excess spikes for right, which is itself proportional to the accumulated logLR that this excess was observed because motion was in fact rightward (***Gold and Shadlen, 2001**; **Shadlen et al., 2006***). For the wider class of decisions, such difference variables are elusive, whereas the possibility of associating a sample with log-likelihood is a natural dividend of learning and memory (***Yang and Shadlen, 2007**; **Kira et al., 2015**; **Shadlen and Shohamy, 2016***).

The results imply the maintenance of separate decision variables each capable of reconciling decision and choice for the one stimulus dimension. We will consider variations and alternatives below, but there must be separate control of termination and negligible cross talk. Specifically, the state of the accumulated evidence bearing on the direction of motion does not affect the amount of accumulated evidence required to reach a decision about color dominance, and the same can be said about the state of the accumulated evidence about color on the decision about direction of motion. In the model the decision variables, *V*_m_ and *V*_c_, represent the integrated evidence for right (and against left) and for blue (and against yellow). Neural correlates of these 1D processes are known, mainly in the parietal and prefrontal cortex (***Gold and Shadlen, 2007***), although they are organized in pairs: ***R–L, L–K*** (and presumably ***B-Y*** and ***Y-B***). Each of the four processes is the accumulation of positive and negative increments, and each is terminated by an upper bound. Because evidence for *R* and *L* are anticorrelated (likewise for ***B*** and ***Y***), the pair of opposing processes is approximated by 1D drift-diffusion to symmetric upper and lower terminating bounds. All model-fits adopt this approximation.

An alternative formulation, which we term target-wise integration, would accumulate evidence for the pair of features associated with each choice target (e.g., ***RB, RY, LB, BY***). If such mechanism were to terminate when the total accumulation reaches a threshold, it would predict a type of choice-interference such that sensitivity to motion, say, would be impaired when the color strength was high, because the decision time is shortened by the stronger stimulus. We have not pursued all variants of target-wise integration, but critically, the bimanual experiment demonstrates that the double decision comprises two terminating events. There may well be neurons that represent the target-wise accumulation of evidence, but they would require additional mechanisms that process color and motion until the first decision terminates. At that time, a threshold could be applied to the target-wise accumulators at a level equal to the sum of the color and motion thresholds, and only the unfinished dimension contributes to the decision. A solution of this type seems a likely possibility in areas of the brain that represent the decision variable as an evolving plan of action.

We find it useful to characterize integration as the implementation of a sequence of instructions to increment and decrement persistent activity in cortical areas that represent the decision variables. In Fig. 8, the instructed change is realized by simple 1st order dynamics chosen to approximate neural responses from area LIP. The implementation is merely phenomenological, but it jibes with emerging ideas in theoretical neuroscience that characterize computation as a change in circuit configuration to establish stable states and dynamics (***Remington et al., 2018***). For decision making, it replaces the requirement for continuous integration, with the realization of instructions as if drawn from a memory stage. This characterization also extends to the buffer.

Recall, the buffer was introduced to explain the observation that a brief pulse of color-motion, acquired in parallel, appears to be incorporated into the decision serially. We characterized the length of the buffer—its storage capacity—using the data from the variable duration experiment Fig. 4, where we equate it with the duration of parallel acquisition. This is reasonable because thereafter, the process is serial. However, this depiction appears to limit the role of the buffer to the beginning of the decision, and it fails to specify how long the information can be held. If there are alternations between color and motion processing before the first process terminates, as shown in Fig. 6, then information might be buffered beyond the initial parallel phase. As shown in Fig. 8C, during alternation a sample might be held for at least 2*τ*_ins_—that is, the time it takes the cleared sample to instruct the appropriate decision process and the time the bottleneck is in play while the other dimension performs its update. If the alternations are less frequent, the buffer might need to hold information longer, and if there is only one transition, then the buffer might be expected hold a sample of information for the duration of the entire first decision. There is presumably a limit on how long a sample can be stored, but studies of visual iconic memory suggest that a sample of evidence might be buffered for ~500 ms (***Sperling, 1960**; **Gegenfurtner and Kiper, 1992***).

We conceive of the buffer residing between the cortical areas that represent the filtered evidence and other cortical circuits that represent the decision variables. Notice that the operations depicted in Fig. 8 assign two duties to the buffer: (i) storage of a sample of evidence while the bottleneck precludes updating the associated decision variable and (ii) conversion of the sample into an instruction to update a decision variable by Δ*V*. These duties could be carried out by different circuits. An appealing candidate for both operations is the striatum. The striatum receives input from the extrastriate visual cortex (***Ding and Gold, 2012a***), and it is known to play a role in connecting value to action selection (***Hikosaka et al., 2014***) as well as working memory (***Akhlaghpour et al., 2016***). In the context of our results, we would characterize the operation as follows. A sample of filtered evidence, represented by the firing rates of neurons in extrastriate cortex (e.g., areas MT/MST) leads to a change in the state of a striatal circuit, such that its reactivation transmits the Δ*V* instruction to the cortical areas that represent the decision variables, and this takes time (*τ*_ins_). On this view, the bottleneck is the striato-thalamo-cortical pathway. There has been an observation of the bottleneck in a split-brain patient, supporting such a subcortical bottleneck at least in certain instances (***Pashler et al., 1994***).

A second possibility is that the buffered evidence is stored in visual cortical association areas, especially areas with persistent representations. For example, it has been suggested that short term visual iconic memory is supported by the slowly decaying spike rates of neurons in area V2 (***O’Herron and von der Heydt, 2009***) and the anterior superior temporal sulcus (STSa) (***Keysers et al., 2005***). This would place the bottleneck between extrastriate cortex and the parietal and prefrontal areas that represent the decision variable (see also ***Marti et al. 2012***). This possibility does not provide an explanation for why communication between these areas would impose a substantial delay (e.g., *τ*_ins_).

A third possibility would identify the buffer with control circuitry within the very cortical areas that represent the decision variables. This might seem far-fetched but there is evidence for such an operation in the premotor cortex of mice, where it underlies the implementation of the logical ‘exclusive or’(XOR) operation (***Wu et al., 2020***). In that case the bottleneck would be intracortical. It would correspond to the implementation of a circuit state from its “silent” representation—that is, in cellular and subcellular (e.g., synaptic) states rather than persistent spike activity. The bottleneck is the conversion from this state to the establishment of the spiking dynamics that instantiate the Δ*V* instruction. This might resemble the recall of an associative memory, which must facilitate the establishment of cortical persistent activity in a state suitable for computation, be it for further updating or comparison to a criterion. The three possibilities are not mutually exclusive; nor are they exhaustive. In any case, the instigating event is the clearance of the bottleneck, signaled by the circuit that receives the Δ*V* instruction.

This brings us to the bottleneck itself. Up to now we have alluded to the bottleneck as a temporary obstruction to color or motion processing, but the bottleneck itself does not add time. It is the instructive step that takes time (*τ*_ins_). This step comprises the conversion of a sample of evidence to a Δ*V* instruction and its transmission to a cortical circuit. Indeed, the same delay is encountered in simpler decisions. For example, in the 1D random dot motion task, the incorporation of evidence into the neural representation of the decision variable is first evident ~180 ms after direction neurons in area MT exhibit direction selective responses (***De Lafuente et al., 2015**; **Ding and Gold, 2012b**; **Kim and Shadlen, 1999***) and this delay holds for perturbations of the stimulus throughout decision formation (***Huk and Shadlen, 2005***). This is too long to be explained by synaptic latencies. It implies either a complex routing through intermediate structures or more sophisticated processing that serves to facilitate the linkage and/or the conversion of the sample to an instruction suitable for establishing the cortical dynamics that ultimately realize the Δ*V* instruction. The delay corresponds to the sum, *τ*_s_ + *τ*_ins_, (circles 1 and 2 in Fig. 8).

Decision variables are represented in the persistent activity of neurons in the parietal and prefrontal cortex of primates. Such persistent activity is associated with working memory, attention and planning. This functional localization conforms to the notion of a “response selection” bottleneck hypothesized by Harold Pashler to explain dual task interference (***Pashler, 1994***), in particular a phenomenon known as the psychological refractory period (PRP): the prolonged latency of the second of two adjacent decisions without an effect on accuracy. In his and our formulation, it reflects a limitation that restricts the flow of information to affect higher processes such as decision-making and short-term working memory. On initial consideration, there is no obvious reason why the formation of working memory should necessitate a bottleneck. If acquisition can be parallel, why not working memory? equivalently, the formation of a provisional plan or intention.

Framed in the language of decision-making, seriality arises as a consequence of limited connectivity between the brain’s evidence acquisition systems—sensory, memory, and emotion—and the systems that represent information in an intentional frame of reference, that is, as provisional affordances. Any possible intention might be informed by a variety of sources of evidence, which may be acquired in parallel but from different locations in the brain. The brain lacks the anatomy to support independent connections from all sources of evidence to all possible intentions—that is, the circuits that represent them. Instead the communication must share connections, and this invites some form of time-slice multiplexing. It is not possible for every source of evidence to communicate with the circuits that form decisions at the same time. For some dedicated operations, it is likely that many sources of “evidence” do converge on the same intentional circuitry (e.g., escape response; ***Evans et al. 2018**; **Lee et al. 2020***), and the tracts can be established through development. But, for flexible cognitive systems that learn and solve problems, the connections between evidence and intention must be multipotent and malleable, since connecting *N* sources of evidence and ***M*** intentions will need at least ***W × M*** wires if they are connected exhaustively, whereas if they are routed centrally, it will only need ***N + M***. This solution necessitates some type of multiplexing (***Zylberberg et al., 2010***)).

We suspect that the constraints leading to serial processing in the color-motion task also apply to other decisions and cognitive functions. For example, deciding between two familiar food items can take a surprisingly long time when those items are valued similarly. This holds when the items are both highly valued or both undesired or both of moderate value. Like decisions about the direction of random dot motion, there is a lawful relationship between the RT to choose an item and the likelihood that the preference is consistent with one’s previously stated value (***Krajbich et al., 2010**; **Krajbich and Rangel, 2011***). Like the choice-RT accompanying 1D motion (or color) decisions, the relationship suggests that some type of process like noisy evidence accumulation—or more generally, sequential sampling with optional stopping—reconciles choice and decision time. However, such expressions of preference differ from perceptual decisions in two important ways. First, there is no objectively correct response, only consistency with the sign of the inequality in the decision-maker’s valuations of the individual items, which are ascertained before the experiment. Second, the food items are not shown as a movie and there is no uncertainty about their identity. Therefore it is not clear what gives rise to independent samples of evidence. Bakkour et al (2019) showed that the samples are likely to arise through constructive processes using hippocampal memory systems. This begs the question why this process would unfold in time like a movie of random dots. An attractive idea is that the use of memory guided valuation—in particular the step to enable it to affect a decision variable—encounters a bottleneck. Even if memories could be retrieved in parallel, they would require buffering and serial updates of the decision variable (***Shadlen and Shohamy, 2016***).

While it is unsurprising that a movie of random dots supplies evidence to be incorporated serially toward a decision, it is shocking that two samples of evidence, supplied simultaneously by the same dots and acquired through parallel sensory channels, do not support simultaneous decisions. In the experiments that require prolonged viewing, non-simultaneity manifests in serial time-multiplexed alternation of the decision processes and the failure to incorporate all information in the stimulus stream into one or both decisions. In a free response design the decision maker compensates by acquiring more evidence, so the interference is not apparent in the accuracy of the perceptual choice. However, if such compensation is precluded by the experimenter (variable duration experiment), the failure to incorporate information can affect accuracy too. That this bottleneck arises despite parallel acquisition of color and motion (or motion from two locations), whether we use one or two effectors to express the decision, and whether we decide between 2 × 2 conjunctions or two categories (same/different) suggests that the bottleneck is pervasive. In addition to the PRP, we suspect that it plays a role in other psychological phenomena, such as post-stimulus masking, iconic memory, the attentional blink, rapid sequential visual processing, and conjunction search. These phenomena represent forms of sequential interference and all can be stated as challenges to the brain’s routing system (***Zylberberg et al., 2010***).

On the other hand, one must wonder if the brain can ever take advantage of parallel acquisition to perform cognitive functions in parallel. It certainly seems so to a musician using their feet and hands to convey time and sonority on a piano or counter rhythms on a drum kit. Yet the time scales of alternation discussed in this paper are on the order of 10 Hz. It seems possible that we achieve parallel processing despite the bottleneck by enhancing signal-processing at the filter stage before the bottleneck and by grouping (or chunking) processes after the bottleneck in higher order controllers of movement and strategy. For example, face selective neurons compute conjunctions of features in less than 100 ms. This is just one example of the sophisticated properties of association sensory neurons in the extrastriate visual cortex, and analogous operations are presumed to occur in secondary somatosensory cortex and belt regions of the auditory cortex. Similarly, complex movement sequences and the rules to coordinate them may be specified in premotor cortex or at the level of the controller. If so, then the only way to overcome the bottleneck is to develop the expertise of the reader or the musician/athlete, leaving most of flexible cognition to negotiate the bottleneck between the acquisition of information and its incorporation into representations that support states of knowledge: decisions, working memory, plans of action. It is the price the brain pays to use its senses (and memory) to bear on a plethora of possible intentions, despite its limited connectivity. The payment is in time, but in another sense, it is time well spent, for without seriality of thought there is no contour to our experiences, no appreciation of cause and consequence, no meaning or narrative.

## Acknowledgments

We thank Daphna Shohamy and Mariano Sigman for contributions to the theoretical underpinnings of our study, and we thank Stanislas Dehaene, Gabriel Stine, Naomi Odean, and Aniruddha Das for comments on an earlier draft of the manuscript.

## Methods

### Participants

Thirteen participants (5 male and 8 female, age 23–40, median = 26, IQR = 25-32, mean = 28.3, SD = 5.74) provided written informed consent and took part in the study. All participants had normal or corrected-to-normal vision and were naïve about the hypotheses of the experiment. The study was approved by the local ethics committee (Institutional Review Board of Columbia University Medical Center).

### Apparatus

Visual stimuli were displayed on high resolution CRT monitors with 75 Hz screen refresh rate. The experiments were conducted in two labs. Table 1 lists the display parameters used in the four experiments. In the eye-tracking experiments, a head- and chin-rest was used, and eye position was monitored at 1 kHz using an Eyelink 1000 device (SR Research Ltd., Mississauga, Ontario, Canada). In the reaching task participants used robotic handles (vBots, ***Howard et al. 2009***) to indicate their choices, and movement trajectories were recorded at 1 kHz. The experiments were run using Matlab and Psychtoolbox (***Brainard, 1997***) and for the online experiments jsPsych (***De Leeuw, 2015***).

**Table 1.** Experimental parameters.

|  | choice-RT (eye) | brief duration (eye) | variable duration (eye) | choice-RT (arm) | binary choice-RT |
| --- | --- | --- | --- | --- | --- |
| Dot density (dots deg<sup>-2</sup> s<sup>-1</sup>) | 15.3 | 15.3 | 16 | 16 | 16 |
| Dot speed (deg/s) | 1.67 | 1.67 | 5 | 5 | 5 |
| Central fixation diameter (deg) | 0.4 gray circle | 0.4 gray circle | 0.6 red cross & bullseye | 0.6 red cross & bullseye | 0.6 red cross & bullseye |
| Random delay (s) | 0.1-0.5 | 0.1-0.5 | 0.5-0.8 | 0.5-0.8 | 0.4-0.8 |
| Choice target diameter (deg) | 0.4 | 0.4 | 1.2 | N/A | N/A |
| Target spacing (deg) | 6 | 6 | 15 | N/A | N/A |
| Movement initiation | gaze > 2.5° | gaze > 2.5° | gaze > 3° | hand > 1 cm | key press |
| CRT | Vision Master 1451 | Vision Master 1451 | Sony CRT CPD-G420S | Dell CRT P1110 | N/A |
| Resolution (pixels) | 1400 × 1050 | 1400 × 1050 | 1280 × 1024 | 1280 × 1024 | S7: 1280 × 720; S12: 1440 × 900 |
| Pixels per degree | 39.6 | 39.6 | 32.7 | 24.3 | S7: 40.94; S12: 33.45 |
| Viewing distance (cm) | 55 | 55 | 50 | 38 | S7: 54; S12: 38 |
| Cyan *cd/m<sup>2</sup>* [M(SD)] | N/A | N/A | 25.20 (0.81) | 12.16 (1.93) | N/A |
| Cyan CIE x/y [M] | N/A | N/A | x = 0.26, y = 0.24 | x = 0.27, y = 0.24 | N/A |
| Yellow *cd/m<sup>2</sup>* [M(SD)] | N/A | N/A | 22.98 (0.05) | 12.68 (1.80) | N/A |
| Yellow CIE x/y [M] | N/A | N/A | x = 0.54, y = 0.38 | x = 0.54, y = 0.38 | N/A |

### Overview of experimental tasks

Participants sat in a semi-dark booth in front of a CRT monitor. They were required to decide the net direction and the dominant color in a patch of dynamic random dots. Individual dots were displayed for a single video frame (1/75 s). Task difficulty for motion was conferred by the probability that in frame *n* + 3 (i.e., Δ*t* = 40 ms), it would be displaced in apparent motion vs. randomly replaced in the aperture. We prepend the probability by plus or minus to indicate the direction, and refer to this signed quantity in units of coherence (coh). For color, task difficulty was conferred by the probability that a dot would be colored blue or yellow on each frame. We refer to the signed quantity, 2(*p*_blue_ – 0.5), as the color coherence. Both coherences share the range {−1,1}. Throughout, we use positive coherence for rightward and blue dominant stimuli. The coherences were stationary during a trial but randomized independently across trials. A calibration procedure was used to match the luminance of the blue and yellow for each participant (see below). For the first experiment (choice-reaction time, participants S1-S3) the color of the dots in the first three frames of a trial was balanced to give no net color information. The procedure was intended to match the state of the motion stimulus which is effectively zero-coherence until the fourth video frame. Subsequent experience demonstrated that this procedure was unnecessary, and we discontinued this practice for the other experiments.

We conducted two types of tasks, a double-decisions (2D) in which both the dominant color and motion direction were reported on each trial, or single-decisions (1D) in which only the dominant color or net motion direction were reported (as in ***Mante et al. 2013***). For 1D experiments the “irrelevant” dimension was varied from trial to trial just as in the 2D task. Variations on this basic design are described in the following sections. We first describe the choice-reaction time task (eye) and then the differences for the other experiments.

For each experiment, the sample size was determined based on prior psychophysics studies with within-subject designs (***Palmer et al., 2005**; **Resulaj et al., 2009**; **Zylberberg et al., 2012**; **Kiani et al., 2014***). Furthermore, trial numbers were chosen such that the number of trials within each dimension given the other dimension’s strength were similar to prior studies (e.g., ***Kang et al. 2017***). We recruited three participants for the first and second experiment (Choice-reaction time task and short duration, eye). For the remaining experiments we recruited 2–8 participants. A larger number was necessary for the arm experiments because fewer trials per hour are acquired and the effort is greater. Unless otherwise stated, participants were randomly allocated to experiments.

### Choice-reaction time task (eye)

Three participants (1 male and 2 female, aged 25–40) performed the task in which they could view the random dots until ready with a response (Fig. 1a). Participants were required to fixate a central spot for 0.5 s to initiate a trial. In the main task (2D), four choice targets appeared at four corners of the display, evenly spaced from each other and the same distance from the fixation spot. The top two targets were colored yellow and the bottom two blue, consistent with the color choices they indicate. For example, to report rightward motion and yellow color, the participant would saccade to the top right target, which was yellow. After a random delay, a patch of dynamic random dots appeared which were restricted between invisible circles of diameter 1 and 5° centered on the fixation spot. The random dots were extinguished when the participant initiated the choice response. Participants were required to respond within 5 seconds of the stimulus onset. Trials in which no response was initiated and those aborted by breaking fixation were repeated at a later time in the experiment. At the end of each trial, the correct target was marked on the screen, and auditory feedback was provided when both dimensions were judged correct.

Participants performed three trial types: color-only, motion-only and color-motion (i.e., double) decisions. For color-motion trials four targets were displayed as in the experiments described above. For motion-only trials two white targets were shown to the left and right of the stimulus, respectively. For the color-only task, one blue and one yellow target were presented above and below the center of the screen, respectively. Participants performed the three trial types in separate 13-min blocks in a random order, in 24–49 blocks over 11–17 days (4775–10973 trials). For the 2D task, 5 strengths (or 9 signed coherences including 0) were used on both dimensions (see Table 2). The set of non-zero motion strengths was doubled for one participant (S1) because they failed to achieve >90% correct at coh=0.256 during training. Likewise, the range of color strengths was doubled for two participants (S1 and S3). For the 1D task, two of the strengths were not used for the irrelevant dimension.

**Table 2.**
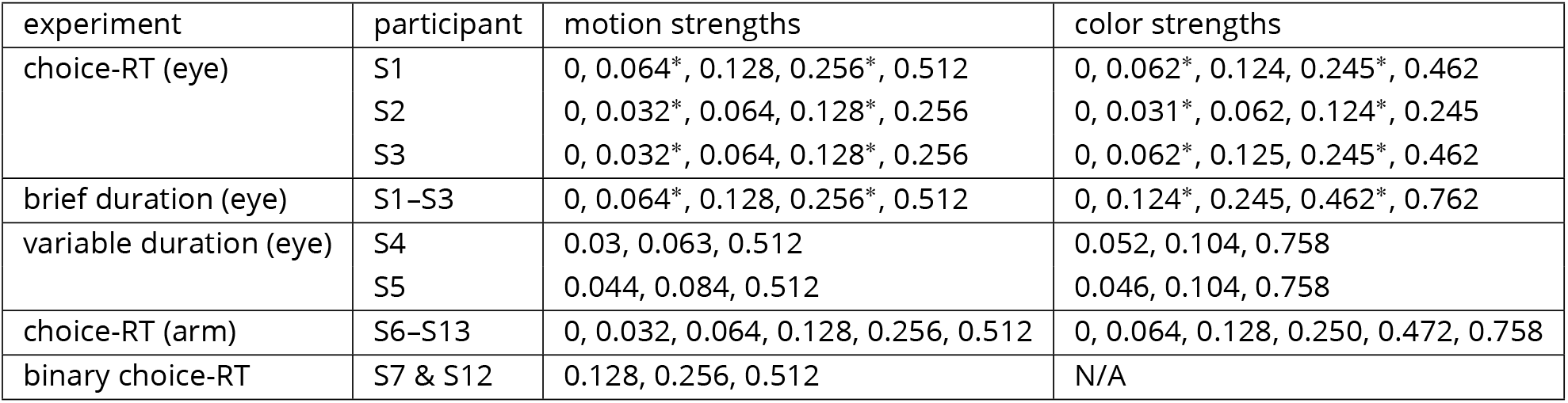
Motion and color strength parameters. For 2D trials all combinations of motion and color strengths were used. For 1D trials all strengths were used for the dimension that informed the decision but some strengths (*) were omitted for the other dimension.

| experiment | participant | motion strengths | color strengths |
| --- | --- | --- | --- |
| choice-RT (eye) | S1 | 0, 0.064*, 0.128, 0.256*, 0.512 | 0, 0.062*, 0.124, 0.245*, 0.462 |
|  | S2 | 0, 0.032*, 0.064, 0.128*, 0.256 | 0, 0.031*, 0.062, 0.124*, 0.245 |
|  | S3 | 0, 0.032*, 0.064, 0.128*, 0.256 | 0, 0.062*, 0.125, 0.245*, 0.462 |
| brief duration (eye) | S1–S3 | 0, 0.064*, 0.128, 0.256*, 0.512 | 0, 0.124*, 0.245, 0.462*, 0.762 |
| variable duration (eye) | S4 | 0.03, 0.063, 0.512 | 0.052, 0.104, 0.758 |
|  | S5 | 0.044, 0.084, 0.512 | 0.046, 0.104, 0.758 |
| choice-RT (arm) | S6–S13 | 0, 0.032, 0.064, 0.128, 0.256, 0.512 | 0, 0.064, 0.128, 0.250, 0.472, 0.758 |
| binary choice-RT | S7 & S12 | 0.128, 0.256, 0.512 | N/A |

**Table 3.** Parameter values for the best-fitting switching model.

| participant ID | $\tau_{\Delta}$ (s) | $p_{\text{motion-1st}}$ | $T_{\text{nd}}^{\text{1st}}$ (s) |
| --- | --- | --- | --- |
| 6 | 1.93 | 0.52 | 0.51 |
| 7 | 1.6 | 0.15 | 0.59 |
| 8 | 4.49 | 0.86 | 0.38 |
| 9 | 1.33 | 0.87 | 0.44 |
| 10 | 0.15 | 0.14 | 0.38 |
| 11 | 0.22 | 0.89 | 0.7 |
| 12 | 0.1 | 0.79 | 0.52 |
| 13 | 1.13 | 0.85 | 0.69 |

#### Minimum-motion procedure

Prior to the experiment, we calibrated the two colors (yellow and blue) to be equiluminant using the minimum-motion procedure (***Cavanagh et al., 1987***). Two vertical sinusoidal gratings with a spatial frequency of 1.25 cyc/deg were shown with a temporal frequency of 6.25 Hz. The first grating had alternating yellow and blue, and the second grating had alternating light and dark green, and they were arranged in a way that if yellow were brighter than blue, the gratings appear to move in one direction (e.g., left), and vice versa. The participant adjusted the luminance of yellow until they did not see net motion, starting from a random luminance value. After 24 trials, the mean luminance of the yellow was computed and used throughout the experiment for the participant.

#### Training sessions

Participants completed 11–13 training blocks (13 minutes, 200 trials) over 4–7 days, beginning with either an easy motion or color 1D task (counterbalanced across participants) and with viewing durations controlled by the experimenter. The incorporation of weaker stimulus strengths and the range of stimulus durations were adjusted progressively. Transitions to the next level were made if the participant met fixation requirements and achieved >90% accuracy on the strongest coherence. The aim was to identify four levels of motion strength ≥ 0.032 and four levels of color strength ≥ 0.031 in octaves steps such that the strongest level (8 times the lowest logit) supported >90% accuracy. We then changed from variable duration to the reaction time version of the 1D task, again ensuring that the range of difficulties led to at least 90% accuracy for the easiest condition. We then repeated these steps for the other stimulus dimension before introducing the 2D choice-RT task. They received a session of practice to gain familiarity with the 4-choice design. For participants S1 and S3, we made a final adjustment of the difficulty levels. The stimulus strengths were then fixed for all test sessions (Table 2).

### Brief duration task (eye)

The same participants from the choice-reaction time task then performed a task that was identical except that the dynamic random dots turned off after 120 ms from the onset. Participants were free to respond after the offset of the dynamic random dots. The “RT” in this task was measured as the time between the onset of the stimulus and the response (the time the gaze left the center of the screen).

Participants completed a total of 35–43 test blocks that each lasted 13 minutes (7309–7745 trials over 12–19 days). The stimulus strengths used are listed in Table 2.

### Variable duration task (eye)

Two participants (2 female, aged 26 and 32; both right-handed) participated and completed a total of 12-26 test sessions that each lasted between 1-2h.

After a training phase (see below) the task alternated between blocks of 72-144 trials where participants either performed the 2D variable duration task, a 1D variable duration task or 2D choice-reaction time task. The majority of blocks were 2D variable duration (total of 11,808 trials). Ten fixed stimulus durations ranging from 120-1200 ms (in steps of 120 ms) were presented in pseudo-random order. Warning messages were displayed if participants initiated an eye movement before the end of the stimulus (“too early!”) or if a movement was not initiated within 5 sec of stimulus offset (“too slow!”). In both cases, the trial was aborted and repeated at a later, randomly determined, trial within the same block.

Only three levels of difficulty were used for each dimension: one easy and two difficult coherence levels. The easy coherence level was 0.512 for motion and 0.758 for color. The two difficult coherence levels were adjusted individually in order to match color and motion performance. Specifically, low coherences for each dimension were chosen to yield 65% and 80% accuracy on each dimension, respectively, based on participants’ performance in the final two training sessions (double-decision RT). All low-coherence levels were < 0.1 for both participants. All 3 × 3 combinations of motion × color were presented. However, since the main model predictions are based on a comparison of trials with hard-hard vs. hard-easy combinations, easy-easy combinations were only presented in 2.4% of trials. All other coherence combinations were presented with equal frequency and counter-balanced within each stimulus duration. Participants also completed 2,160 trials each of motion-only and color-only trials and 1,296 trials of the 2D choice-RT task which were included to ensure that they maintained appropriate speed-accuracy trade-offs throughout the experiment.

#### Training sessions

Participants first completed 6-9 training sessions. In the first 2 sessions, they were trained on a variable duration task where stimulus durations were drawn randomly from a truncated exponential distribution ranging between 500-2000 ms (session 1) or 100-1600 ms (session 2). Participants first completed 1D-motion and 1D-color tasks in separate blocks, followed by the 2D task. In the remaining training sessions, participants mainly performed a 2D RT task until they reached stable performance (at least 60% accuracy on the 2nd coherence level for both decision dimensions, with little to no changes in choice performance or RTs over blocks). Occasionally, additional 1D blocks were introduced in order to obtain similar performance levels for motion and color judgments. Throughout training, all 6 coherence levels for motion {0,0.032,0.064,0.128,0.256,0.512} and color {0,0.064,0.128,0.250,0.472,0.758}, and all their possible combinations, were presented.

#### Isoluminance calibration

At the start of the experiment, participants completed a flicker fusion procedure to match luminance of yellow and blue. A square (4.9 × 4.9°) was presented in the center of the screen. The color of the square flickered at 37.5 Hz between cyan and yellow. For efficiency we only explored values [***R G B***] = [0 *x x*] and [***R G B***] = [*y y* 0], for cyan and yellow, respectively, where 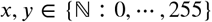. Participants pressed the left or right arrow key to minimize the perceived flicker. One key changed *x* and *y* by +1 and −1, respectively, and the other key had opposite effect. Participants pressed the space bar to signal the subjective point of minimal flicker. This procedure was repeated 10 times, each time starting with new initial values [*x y*], chosen pseudo-randomly, such that either yellow or blue was dark while the other color was bright (counter-balanced across trials). The precise initial values were chosen to be equidistant from 225 and were between 195-200 for the darker color and 250-255 for the brighter color (e.g., blue would start at [0 197 197] and yellow would start at [253 253 0]). This ensured sufficient contrast to induce a flicker at the start of each trial. The averages across the 10 trials were adopted as the isoluminant setting for the participant. After the procedure, participants were presented with a single trial with the obtained color values and were asked to report if they perceived a flicker. If they did, the procedure was repeated. The same calibration procedure was also used for the next experiment.

#### Choice-reaction time task (arm)

Twelve right-handed participants were initially recruited for the experiment. After training, 8 participants were selected for the actual experimental sessions based on their overall performance. Participants completed two test sessions with a unimanual version of the task and two test sessions with a bimanual version (order counterbalanced across participants). In each experimental session, all 6 color × 6 motion strengths combinations were presented – that is {0,0.032,0.064,0.128,0.256,0.512} × {0,0.064,0.128,0.25,0.472,0.758} – pseudo-randomly in 12 blocks of 96 trials each (total of 1152 trials). The order of unimanual and bimanual sessions was counterbalanced across participants.

Unlike the eye experiments, no choice targets were present on the screen. Instead, there were arrow icons that indicated the mapping of color and motion to forward/backward (appropriately colored) and left/right directions of the hand (Fig. 5A). The mapping of blue/yellow to bottom/top target locations was counterbalanced across participants. The movements themselves were restricted to virtual channels in the plane. In the unimanual task, participants moved a single robotic handle with either their left or right hand (counterbalanced across each half of a session) in one of the 4 diagonal target directions (2 color × 2 motion; as in the other experiments). In the bimanual task, participants used two separate robotic handles to move their left and right hand in a left/right (motion judgments) and forward/backward (color judgments) direction, respectively (hand assignment counterbalanced across participants). Feedback about the hand position(s) was provided by two black bars on top of the arrow icons (for clarity shown as grey in Fig. 5A). Participants were instructed to move each bar in the chosen direction until their hand(s) reached a virtual ‘wall’ at the end of the channel, at which point their decisions were registered. Movement distances between starting positions and target locations were identical in the uni- and bimanual task (5 cm). On 2D trials the random dots were extinguished when both decisions were indicated, that is when the hand left the home position in the unimanual task and when both hands had left the home position in the bimanual task. Warning messages were presented if participants initiated a response before stimulus onset (“too early”) or when RTs exceeded 5 sec (“too slow!”). In both cases, the trial was aborted and was repeated at a later, randomly determined, trial within the same block.

Once participants indicated their decision, green/red frames were presented around the response arrows to indicate correct/incorrect choices separately for each decision dimension. If both decisions were correct, additional auditory feedback was provided (700 Hz tone) indicating that participants had won one point. Participants were instructed to maximize points by responding as fast and accurately as possible. At the end of each trial, they received feedback regarding their current rate of rewards (points/min) as well as a graph of their scores in each 2 minute period over the last 10 minutes. To further motivate participants to adopt appropriate speed-accuracy trade-offs, feedback duration was longer for errors than correct responses, hence delaying the onset of the next trial (correct: 1250 ms; error on one dimension: 2000 ms; error on both dimensions: 3000 ms). At the end of the trial the robotic interface actively moved the hand(s) back to the home position(s).

#### Training

All participants completed 3-4 initial training sessions, using the version of the task that they were assigned to first (uni- or bimanual, counterbalanced; see above). In the first two training sessions, participants performed a variable duration task with stimulus durations varying between 500-2000 ms. The third training session introduced the choice-RT design. To train participants to maximally separate their two hands in the bimanual version, the RT training task alternated between easy-motion (motion coherence = 0.512) and easy-color (color coherence = 0.758) blocks, and participants were encouraged to respond as quickly as possible to the easy dimension while taking more time to make a correct choice on the harder dimension. For participants who were first trained on the unimanual version, stimulus coherences were also presented in blocks of easy-motion vs. easy-color to ensure consistency in training across all participants. Participants were invited for the experimental sessions only if their overall rate of warning messages was less than 5% and if their average accuracy was at least 95% on the easy dimension and at least 65% on the 3rd highest coherence level of the harder dimension (motion: 0.064; color: 0.128).

After initial training, participants completed 2 experimental sessions of the task they had been trained on (either uni- or bimanual RT task). They then completed another practice session, in which they were trained on the other version of the task (either bi- or unimanual RT task), before completing 2 final experimental sessions with this version of the task. Experimental sessions only differed in motor implementation of decisions (uni- vs. bimanual), but were otherwise identical, and S-R mappings were kept constant within participants.

### Binary choice-reaction time task

The experiment was conducted remotely during the SARS-CoV-2 pandemic (summer 2020). Two participants who had also completed the uni- and bimanual tasks were recruited for this experiment. Participants completed the task online using a Google Chrome browser on Windows 10 and macOS Catalina (version 10.15.4), respectively. Both participants completed eight separate one hour sessions within a two week time period. The task was programmed in JavaScript and jsPsych (***De Leeuw, 2015***).

During the task, two random dot motion patches with rectangular apertures (each 3 × 5°) were presented to the left and right of a red fixation and separated by a central gray bar (2 × 5°) cross Fig. 7A. Motion direction (up/down) and coherence ({0.128, 0.256, 0.512}, referred to as low, medium and high) of the two stimuli were independent of each other. The six unique coherence combinations were presented with equal frequency and in randomized order. The stimuli directions and allocation to the left vs. right side of the screen were counterbalanced. Participants had to judge whether the dominant motion directions of the two stimuli were the same or different and indicate their choice by pressing the F or J key with their left/right index finger, respectively, when ready. The response mapping was counterbalanced across the two participants and was shown at the bottom of the screen throughout the task. Visual feedback was provided at the end of each trial. For correct responses, participants won 1 point. After errors and miss trials (too early/late), participants lost 1 point. Miss trials were repeated at a random trial during the same block. Participants were instructed to try and win as many points as possible and they received an extra bonus of one cent for every point they won. Their point score was shown in the corner of the screen throughout the task and additional feedback about percent accuracy was provided at the end of every block.

Participants first completed 3 training sessions after which they completed 4 sessions of the same-different task (3072 trials in total). Finally, participants completed a single session (768 trials) of a 1D task in the random dot motion was restricted to the left or right patch (counterbalanced across trials) and participants had to judge the motion direction (up/down) by pressing the M or K key with their right index/middle finger, respectively.

At the end of each session, participants completed a separate block of 32 trials with 100% coherence stimuli only (sessions 1-7: same-different task; sessions 8: 1D task). Participants were instructed that decisions in this block would be very easy and that they should respond as fast as they could while still being accurate. The reaction times obtained from these blocks (not shown) serve as a check on our estimate of the non-decision time (***Stine et al., 2020***). Participants were instructed to maintain fixation throughout the task. At the end of each session, they provided self-report judgments indicating to what extent they kept fixation during the task on a scale from 1 (“not at all”) to 4 (“always”). The mean and interquartile range of the reports were 3.75 and 3.5–4 (combined for the two participants). Prior to the experiment, participants completed a virtual chinrest procedure in order to estimate viewing distance and calibrate the display in terms of viewing angle (***Li et al., 2020***). This involves first adjusting objects of known size displayed on the screen to match their physical size and then measuring the distance from fixation to the blind spot on the screen (corresponding to around 13.5°).

### Serial and parallel drift diffusion models

Both the serial and parallel models assume that decisions are based on the accumulation of evidence over time. The decision processes for color and motion are described by two independent Wiener processes with drift. The decision variable for one of the dimensions (here motion), evolves according to the sum of a deterministic and a stochastic component:

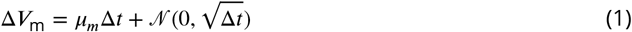

The deterministic term depends on the drift *μ_m_*,

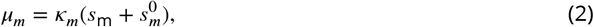

where *s*_m_ is the stimulus motion strength (signed coherence). By convention, *s*_m_ is positive (negative) when the motion is to the right (left). *κ_m_* is a parameter that converts coherence to a signal-to-noise ratio, which we fit to the data. 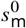 is a bias that allows us to explain, for example, why left and right responses may not be equiprobable even when there is no net motion in either direction. We model the bias term as an offset in the coherence rather than the starting point of the accumulation. This approximates the optimal way of incorporating a bias in drift-diffusion models when there is uncertainty about the reliability of evidence (e.g., the coherence levels vary across trials) (***Hanks et al., 2011**; **Zylberberg et al., 2018***).

The second term of Eq. 1 describes the stochasticity that affects the evolution of the decision variable. It captures the variability introduced by the stimulus and the brain. This variability is modeled as samples from a normal distribution with zero mean. By convention, the standard deviation is 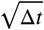, which results in the variance of the decision variable equal to 1 after accumulating evidence for 1 second. This choice does not lead to any loss of generality since for any other value it would be possible to define a new model that has the same behavior in which the variance is 1 and the other parameters are a scaled version of the original ones (***Palmer et al., 2005***).

The accumulation process stops and a decision is made when the accumulated evidence reaches one of two bounds. The choice is ‘rightward’ if the decision terminates at the upper bound, and ‘leftward’ if it terminates at the lower bound. The decision time is the time *T*_m_ that it takes the decision variable to cross the bound. The upper and lower bounds are assumed symmetric with respect to zero. To explain why errors are (often) slower than correct responses, the bounds are allowed to collapse over time. We parameterize the bound as a logistic function with slope *a*_m_. The bound reaches a value of *u_m_*/2 at *t = d_m_* and approaches 0 as *t* → ∞:

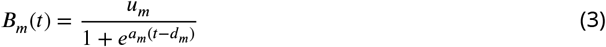

with lower bound simply −*B_m_*(*f*).

Although the previous explanation focused on the motion decision, the same equations describe the decision process for color. We use subscript *c* instead of *m* to refer to the color decision, and adopt the convention that positive (negative) evidence supports the blue (yellow) choice.

Given a set of parameters 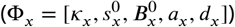, where *x* ∈ {*c, m*}, we can estimate the probability density function for the decisions time *T*_x_, and the two possible choices *R* (right/left for motion and blue/yellow for color). This density function, denoted 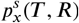, depends on the signed stimulus coherence, *s*. We obtain it by numerically solving the Fokker-Planck equation associated with the Wiener process with drift (***Kiani and Shadlen, 2009***), using the numerical method of Chang & Cooper (1970).

So far, the model description applies to making single decisions (1D) for motion and color. The serial and parallel models are used to explain how combined color-motion decisions (2D) are made. In the serial model the accumulation of evidence at any time can only be for color or motion and therefore the total decision time *T* is the sum of the decision times for motion (*T*_m_) and color (*T*_c_), and the distribution of decision time is given by:

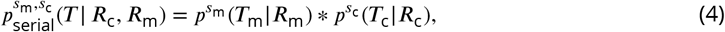

where *R*_c_ and, *R*_m_ are the responses (i.e., choices) for color and motion, respectively, and ∗ denotes convolution.

In contrast in the parallel model both the motion and color are processed simultaneously and, therefore, the decision time is the maximum of either decision time: max(*T*_c_, *T*_m_). We can numerically derive the distribution of decision times from the single-modality distributions by noting that the decision time is equal to *t* if (*i*) motion ended at time *t* and color ended before time t, (*ii*) color ended at time *t* and motion ended before time *t*. Thus,

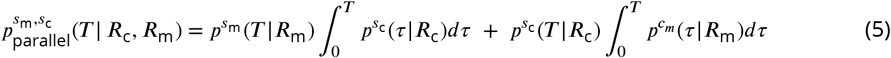

Besides the decision-time, there are sensory, motor and processing delays that contribute to the total response time. We assume that the combined non-decision latencies, *T*_nd_ are normally distributed with a mean of *μ_tnd_* and a standard deviation of *σ_tnd_*. The observed RT distribution for each stimulus condition and choice is then obtained by convolving the distributions of the decision times and the non-decision times, which follows from the assumption that decision and non-decision times are additive and independent.

To avoid over-fitting, our strategy for comparing the serial and parallel models was to fit all parameters using the subset of trials in which one of the two dimensions had maximum strength (Fig. 1b). We used the Bayesian Adaptive Direct Search method (***Acerbi and Ma, 2017***) to search over the space of parameters. The best-fitting parameters are shown in Table 4, for each participant and model type.

**Table 4.**
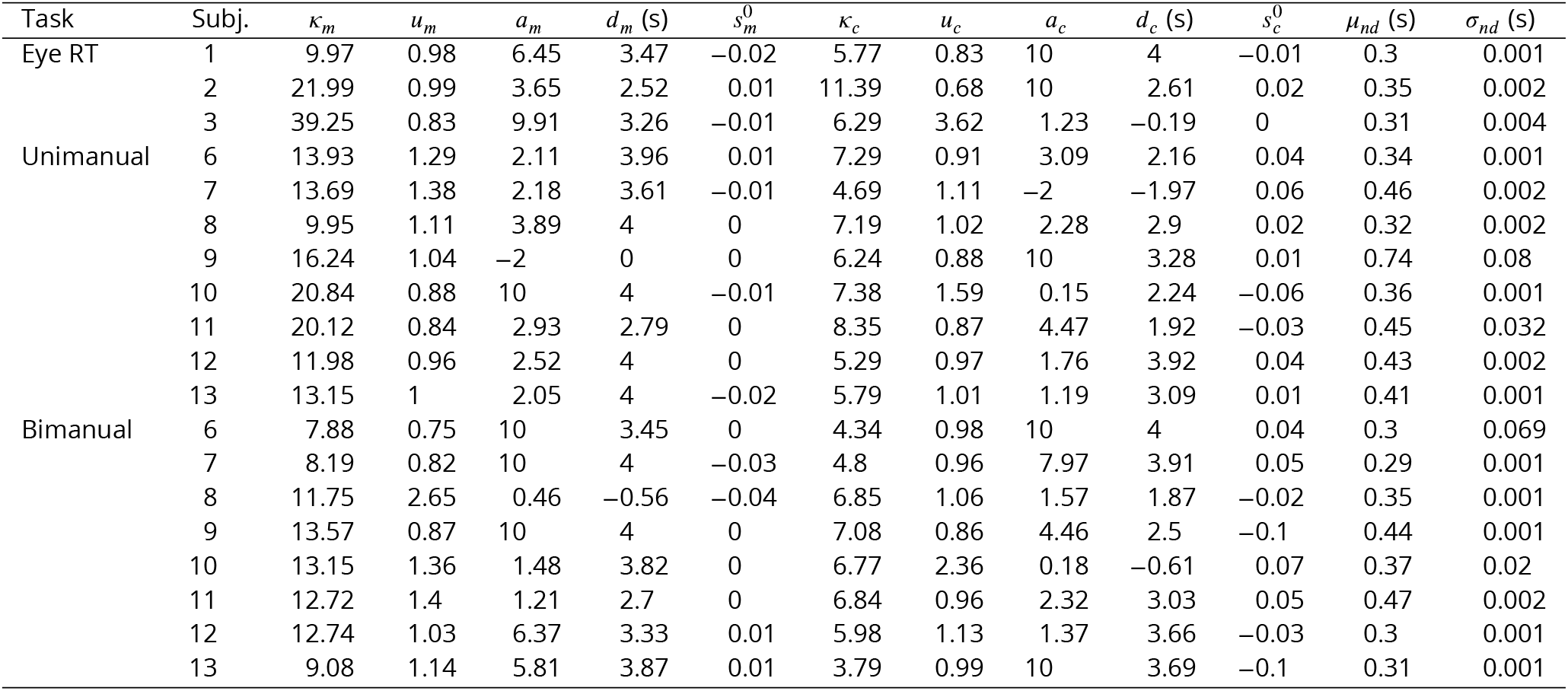
Parameter values for the best-fitting serial model. Note that the rate of collapse parameters *a_m_* and *a_c_* are limited to a maximum of 10 (an almost instantaneous bound collapse) and the time of the start of the collapse *d_m_* and *d_c_* are limited to 4 s.

From the marginal distributions, we predict the choices and response times for all combinations of motion and color coherence, and compare the models by the probability that each one assigns to the data that was not used for fitting. Because the two models have the same number of parameters (*N* = 12: 5 each for Φ_*m*_ and Φ_*c*_, plus 2 for non-decision time), we can directly compare the raw likelihoods (***Figure 2–Figure Supplement 1***).

We conducted a model recovery exercise to verify that our fitting procedure would recover the correct model if the data were generated by either the serial or the parallel model. For each participant and model type (serial/parallel), we generated a synthetic data set with the same number of trials per condition (combination of color and motion coherence) as completed by the participant. The parameters used to generate the synthetic data set were those that best fit the participants’ data (that is, those shown in Table 4). Then we repeated the model comparison (just as we did for the participants’ data) and assessed whether it favored the model that was used to generate the simulated data. ***Figure 2–Figure Supplement 1*** shows that our model comparison procedure can reliably identify the correct model for 37 out of 38 comparisons.

For the binary choice-reaction time task we fit a serial drift diffusion model jointly to the 1D and 2D choices and mean RTs for each participant. The 1D model is simple diffusion to stationary, symmetric bounds, which determine the proportion of up and down choices as a function of motion coherence, as in Eqs. 1 and 2. For the 2D trials we assumed that participants applied the same decision process to each stimulus to determine an up-down choice and that the same-difference response was made by comparing the two decisions. We assumed that sensitivity was the same for the 1D and 2D choices but allowed separate bounds and non-decision times. The application of stationary (i.e., non-collapsing) bounds fails to account for the distribution of RTs and it underestimates the mean RT on errors (***Ratcliff and Rouder, 1998**; **Drugowitsch et al., 2012***). We therefore fit the mean RTs for the correct choices. For the same-different task, we are assuming negligible contribution of double errors (i.e., incorrect direction decisions for both the left and right patch) to the mean RT. The fit maximized the likelihood of the choice assuming binomial error (from the model) and Gaussian error (from the data).

### Comparison of double-decision reaction times under serial and parallel rules

We pursued a second approach to compare serial and parallel integration strategies, focusing specifically on the decision times. Unlike the fits to choice-RT, this method uses each participant’s choices as ground truth. It considers only the distribution of RTs and attempts to account for them under serial and parallel logic. Instead of diffusion models, we estimated the marginal distributions for each 1D decision time with gamma distributions. Specifically, for each motion strength and choice (*s*_m_ & *R*_m_)and each color strength and choice (*s*_m_ & *R*_c_) we modeled the 1D decision time distributions as a gamma distribution (two parameters governing mean and standard deviation). These 1D distributions allowed us to predict the decision time on 2D trials under a serial (additive) and parallel (max) rule. The non-decision times were also modeled as four gamma distributions, one for each combination of the four choices (*R*_m_ & *R*_c_). The reaction time distribution was obtained by convolution of the decision time and non-decision time distribution. Each participant’s data was fit under the serial and parallel model by maximum likelihood (using Matlab fmincon). For robustness, only combinations of strengths and choices with more than 10 trials were included in the fit. The analysis is therefore heavily weighted toward correct trials. Comparison of models was based on log likelihoods of the data given the fitted parameters for each participant.

We validated this method on synthetic data from a parallel and serial simulation and showed that model recovery was accurate (***Figure 2–Figure Supplement 2***).

We also deployed the fit-predict strategy used in Fig. 2, where we estimated the gamma distributions for the 1D decision times and using only the conditions in which one or the other stimulus dimension was at its maximum strength (|*s*|) (***Figure 2–Figure Supplement 5***).

For the binary response task (same/different judgments), a simplified version of this model was used (***Figure 7-Figure Supplement 1***). Only absolute coherence levels of each motion stimulus were considered to fit the marginal gamma distributions. Additionally, only RTs from correct trials were included in this model. Finally, in order to estimate the distribution of *T*_nd_, only a single gamma distribution was fitted.

### Variable duration model

We assume that when the duration of the color-motion stimulus is controlled by the experimenter, the choices are still governed by bounded integration. Thus decisions can terminate (e.g. at time *T*_m_ for motion) before the stimulus duration, *T*_dur_ (***Kiani et al., 2008***). For example in a 1D decision about motion stimulus with strength s_m_, the choice is determined by (1) the distribution of termination times, 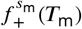 and 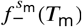, at the positive and negative bounds, respectively, up to *T*_dur_ and (2) the probability that the sign of the unabsorbed *V*_m_(*t = T*_dur_) is of the corresponding sign. For example the probability of rightward decision for a stimulus duration *T*_dur_ is

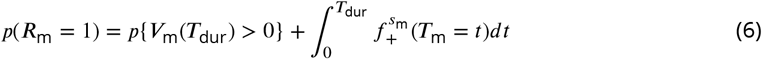

Note that 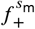 is not a proper density; the total probability at *t = T*_dur_ comprises absorption times at both bounds and the probability of unterminated *V*_m_(*T*_dur_).

To fit the data in Fig. 4 we employ two drift diffusion models, for color and motion, which only interact in the way they access the stream of sensory evidence. This interaction is governed by two parameters, one that determines the amount of time (*T*_buf_) for which processing occurs in parallel before proceeding to a serial processing stage, and the second (*p*_motion-1st_) the probability that motion is prioritized over color during the serial stage. If motion is prioritized on a particular trial, for example, the motion process accumulates evidence in the serial phase until a decision bound is crossed at which point color evidence continues to accumulate. Therefore, if *V_m_* does not reach a decision bound before the sensory stream terminates, no further color evidence is accumulated after the parallel phase.

To model the double-decisions, we used the two 1D processes to specify the duration of the stimulus that was used for motion processing, *T*_m_, and color processing, *T*_c_. On a trial in which motion is prioritized, the time component that contributed to the motion accumulation (*T*_m_) is either the time, *T*_m_, that *V*_m_ reaches a termination bound or *T*_dur_ if it does not reach a bound. These two possibilities bear on the maximum time available for color processing 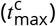:

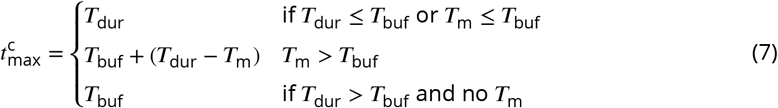

The three conditions in Eq. 7 can be understood intuitively. (1) If the stimulus is shorter than the parallel phase or if motion has terminated in this phase, then the maximum time available for color processing is the full duration of the stimulus. (2) If motion terminates in the serial phase, then the maximum time available for color is the duration of the parallel phase and what time remains of the serial phase after motion has terminated. (3) If motion does not terminate, then color is only processed during the parallel phase. With probability 1 – *p*_motion-1st_, color is prioritized, and the complementary logic holds.

Note that if *T*_buf_ = 0, the model is purely serial with one change from motion to color with probability *p*_motion-1st_ or from color to motion with probability 1 – *p*_motion-1st_ Although realized as a single switch, the model is qualitatively indistinguishable from other alternation schedules that preserve the same competition for processing time. For *T*_dur_ ≤ *T*_buf_, the model is effectively parallel. We fit a parallel model to the data (***Figure 4–Figure Supplement 2***) by fixing *T*_buf_ to the longest duration tested (1.2 s).

Each of the 1D diffusions were modeled similar to those used for the RT task, except for the following minor modifications. (1) We did not include a parameter for nondecision times, because we only modeled choices. (2) We parameterized the bound as an exponential function that is clipped to have a maximum at *u_m_* and start decreasing from *t = g_m_* with a half-life of *d_m_*:

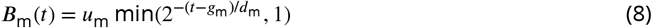

with lower bound simply −*B*_m_(*t*). The same parameterization applies to the color bound (terms with subscript c in Table 5).

**Table 5.** Parameter values for the best-fitting buffer + serial model.

| Task | Subj. | $\kappa_m$ | $u_m$ | $g_m$ (s) | $d_m$ (s) | $s_m^0$ | $\kappa_c$ | $u_c$ | $g_c$ (s) | $d_c$ (s) | $s_c^0$ | $p_{\text{motion-1st}}$ |
| --- | --- | --- | --- | --- | --- | --- | --- | --- | --- | --- | --- | --- |
| VD | 4 | 22.29 | 0.86 | 0.47 | 1.19 | 0.01 | 9.18 | 1.39 | 0.18 | 0.27 | -0.01 | 0.80 |
|  | 5 | 9.60 | 0.99 | 0.68 | 0.40 | 0.01 | 12.11 | 0.74 | 0.11 | 0.11 | 0.04 | 0.96 |

The model was implemented in PyTorch (***Paszke et al., 2019***) with an Adam optimizer (***Kingma and Ba, 2014***) and a modified version of the cyclical learning rate schedule that simply switched back and forth between 0.05 and 0.025 every 25 epochs (***Smith, 2015***). We verified that this procedure reliably recovers the Tbuf (see ***Figure 4–Figure Supplement 1***). Briefly, in Adam, the learning rate gives an approximate upper bound to the change each parameter takes per epoch, and the step size is also adapted for individual parameters based on the running estimates of the first and second moments of the gradient. That is, a high learning rate updates parameters fast and a low learning rate allows better convergence at the expense of speed. We fit the model separately for each *T*_buf_ in steps of 40 ms from 0 to 240 ms and then in steps of 120 ms up to 1200 ms, the longest duration of the stimulus we used. The reported estimate of *T*_buf_ is the sample value with maximum log likelihood. The intervals reported are guided by the observation that choice predictions change little with the buffer duration when the duration is long.

To evaluate the validity of the estimates of buffer capacity (*T*_buf_) shown in Fig. 4, we performed two types of analyses for each participant (***Figure 4–Figure Supplement 1***). The first approximates the specificity, the second the sensitivity of the estimates. (1) We used the parameters of the best fitting diffusion models to the data in Fig. 4 (solid curves; see Table 5) to simulate synthetic data using buffer duration of *T*_buf_ = 80 ms. We fit the synthetic data with models with the buffer capacity fixed to other values (from 0 to 240 ms in steps of 40 ms, and from 240 to 1200 ms in steps of 120 ms). We then compared log likelihood of those fits with that of the 80-ms buffer model, and repeated the simulation 12 times. (2) We used the parameters of the best fitting diffusion models to the data in Fig. 4 to simulate synthetic data using the buffer durations, *T*_buf_ ≠ 80 ms, and compared two fits: with *T*_buf_ = 80 ms or the simulated value.

### Multi-switch model (arm)

In the serial phase of the 2D task, the motion and color processes alternate. Experiments that provide only one response time to report both decisions allow us to estimate the overall prioritization of one stream over the other but not the frequency of alternation. In contrast, the bimanual task provides two response times on each trial. This allows us to estimate the frequency of alternation between stimulus dimensions by fitting a model with multiple switches to the response times of the first decision in the bimanual task.

The fitting was carried out in two steps. First, we fit the serial model described in Eq. 4 to the second response in the bimanual task. The parameters that best fit the data are shown in Table 4. Second, with the serial model parameters fixed, we used three additional parameters to account for the response times to the decision that was reported first. The three parameters are: *τ*_Δ_, controlling the average time between alternations of color and motion; *p*_motion-1st_, the probability of starting with motion; and 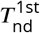 the expectation of the non-decision time for the first response.

The alternations are modeled as a renewal. The intervals are independent and identically distributed (*iid*) as

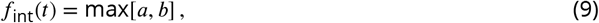

where *a* and *b* are draws from an exponential distribution with mean *τ*_Δ_. The expectation of the interval is

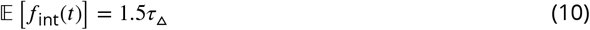

We chose this parameterization so that the distribution of inter-switch intervals has a single peak and the max operation reduced the probability of very short intervals.

Because there is no closed-form solution to the multi-switch model, we used simulations to fit the model parameters to each participants’ data. For fitting, we simulate the model 1,000 times for each unique combination of color and motion strengths. From the simulations, we average the response times for the first decisions split by whether motion or color was reported first, and binned them by both motion strength and color strength. This gives the four groupings in Fig. 6. The parameters were fit to minimize the sum of squared-errors summed over these four groups; in other words, we minimize the sum of the squared errors for the data points shown in Fig. 6. We used this approach rather than maximum likelihood because of the difficulties of reliably estimating the likelihood of the parameters from model simulations for continuous quantities (here, response times) (***van Opheusden et al., 2020***).

### Data Analysis

We used logistic regression to evaluate the influence of task type (single,double) on performance in the short-stimulus duration task (Fig. 3). Separate regression models were fit for the color and motion decisions. The logistic regression model is:

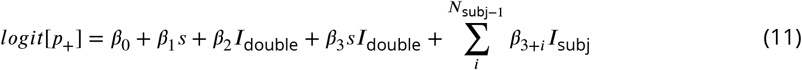

where *p*_+_ is the probability of a positive (‘rightward’ for the motion task, ‘blue’ for the color task) response, *s* is (signed) stimulus strength, *I*_double_ is an indicator variable for task type (single or double), *β*_3_ is an interaction term which indicates how the influence of strength on choice changes in the double task relative to the single task, and *I*_subj_ is an indicator variable that takes a value of 1 if the trial was completed by subject subj and 0 otherwise. The final term with the summation allows for the possibility that different participants had different overall choice biases.

We also used logistic regression to assess whether the strength of one stimulus dimension affected the accuracy of the other decision. Separate regression models were fit for the color and motion decisions. The logistic regression model to assess whether color strength affects motion choice is

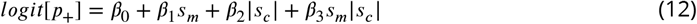

where the *β*_3_ term accommodates the possibility that the color coherence could affect the slope of the logistic function of motion coherence. We used an analogous equation to ask whether motion strength affected color sensitivity. For both logistic regression models (Eq. 11 Eq. 12, to test whether the interaction (*β*_3_) has explanatory power in the model we compared the Bayesian Information Criterion (BIC) for nested regression models with and without the *β*_3_ term. For Eq. 12 data were fit for each participant and the BICs were added.

For the model-free analysis of the time course of the influence of motion and color information on choice Fig. 3, we obtained choice-conditioned averages of the color and motion energies extracted from the random-dot stimuli. Because the stimulus is stochastic, the motion and color energies vary from one trial to another, and even within a trial. We quantified the motion fluctuations by convolving the sequence of random dots presented in each trial with a filter selective to rightward and leftward motion (see details in ***Adelson and Bergen** (**1985**); **Kiani et al.** (**2008**)*). The results of the convolution are combined over space to obtain the motion energy for each direction and as a function of time, and the net motion energy is obtained by subtracting leftward from rightward motion. This time-dependent signal comprises a deterministic component, associated with the motion strength and direction of each trial, and a stochastic component (i.e., each random dot movie uses a unique random sequence of dots). Because only the latter provides information about the time-course of decision formation, we subtracted from the motion energy profile of each trial, the average motion energy associated with the strength and direction of motion of that trial. The motion energy residuals were then averaged across trials, separately for 1D and 2D trials (Fig. 3B).

We performed a similar analysis to extract the color energy from the stimulus. We calculated the difference between the number of blue and yellow dots shown on each video frame. We subtracted the expectation of this difference, given by the color strength of the trial and the predominant color, to obtain the excess of color dots for blue over yellow. These calculations were performed independently for each video frame; the visual system, however, blurs the color information over time. Since we do not know the time constant of this operation, we used an impulse response function that matches the motion filter (Fig. 3B). That is, we convolve the excess of color dots with the temporal impulse response obtained from the motion energy filters. This choice does not affect the conclusions we draw from this analysis – even if we used the unfiltered color residuals, we would still conclude that the same evidence samples were used to form color decisions in 1D and 2D trials.

**Figure 2–Figure supplement 1.**
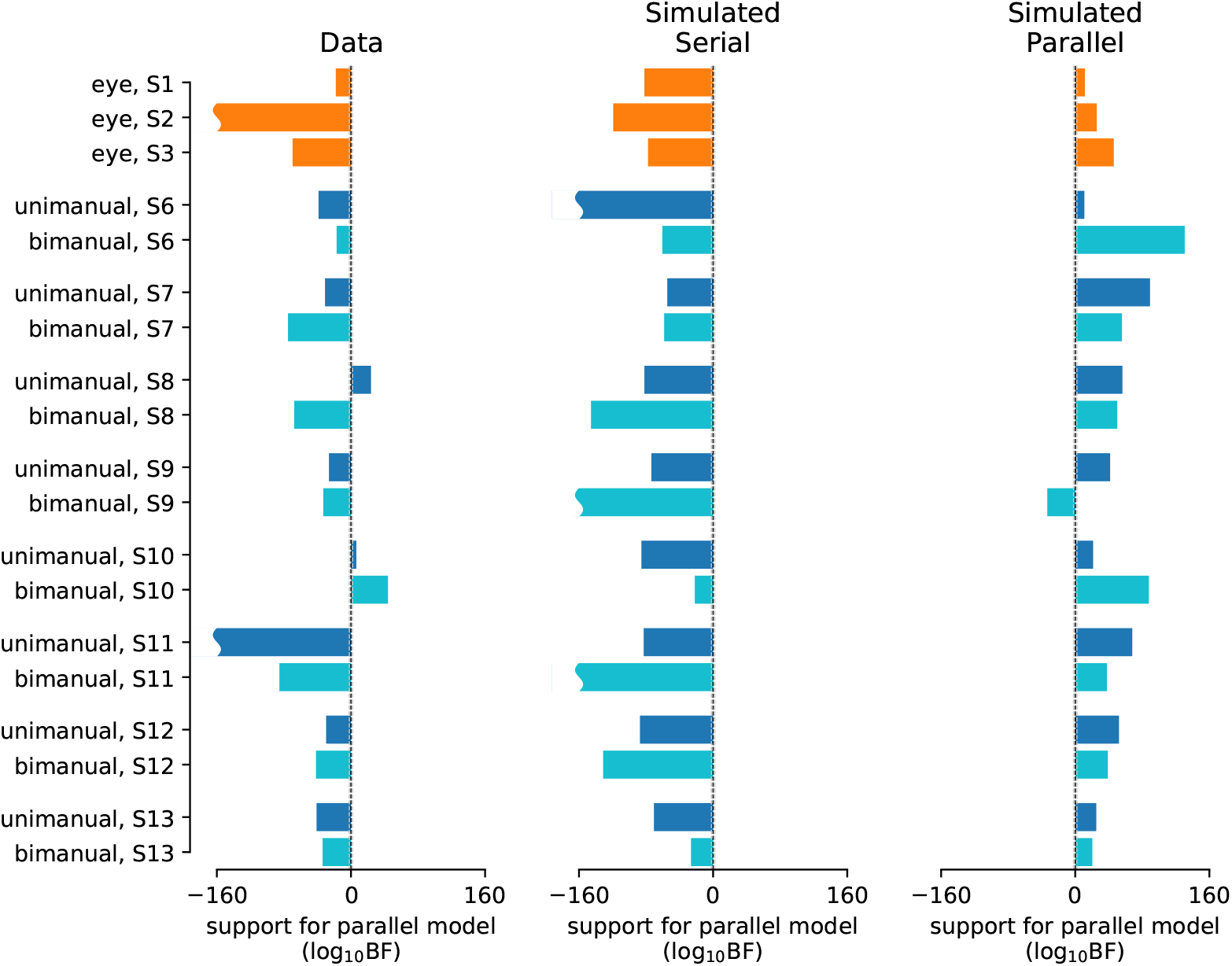
Statistical comparison of the drift diffusion model under serial vs. parallel rules. The analysis focuses on the data and predictions represented by the solid symbols and lines in Fig. 2. *Left*, Difference in log likelihood of the predictions under parallel and serial rules for each participant and condition. Gray vertical dashed lines (close to the midline) show where the Bayes factor is 1/100 and 100 (log_10_BF = −2 and 2; “decisive” evidence in support of the model on that side; ***Kass and Raftery 1995***). Negative and positive values correspond to support for the serial and parallel model, respectively. *Middle and right*, Validation of the method. After fitting, these parameters were used to generate simulated data under the serial (left) and parallel (right) rules. Each dataset was then fit using both the serial and parallel rule. The validation shows that 37/38 simulated datasets were correctly categorized. Average log_10_BF ± SEM are −58±20, −101±14, 44±8 for the data, simulated serial, and simulated parallel, respectively.

**Figure 2–Figure supplement 2.**
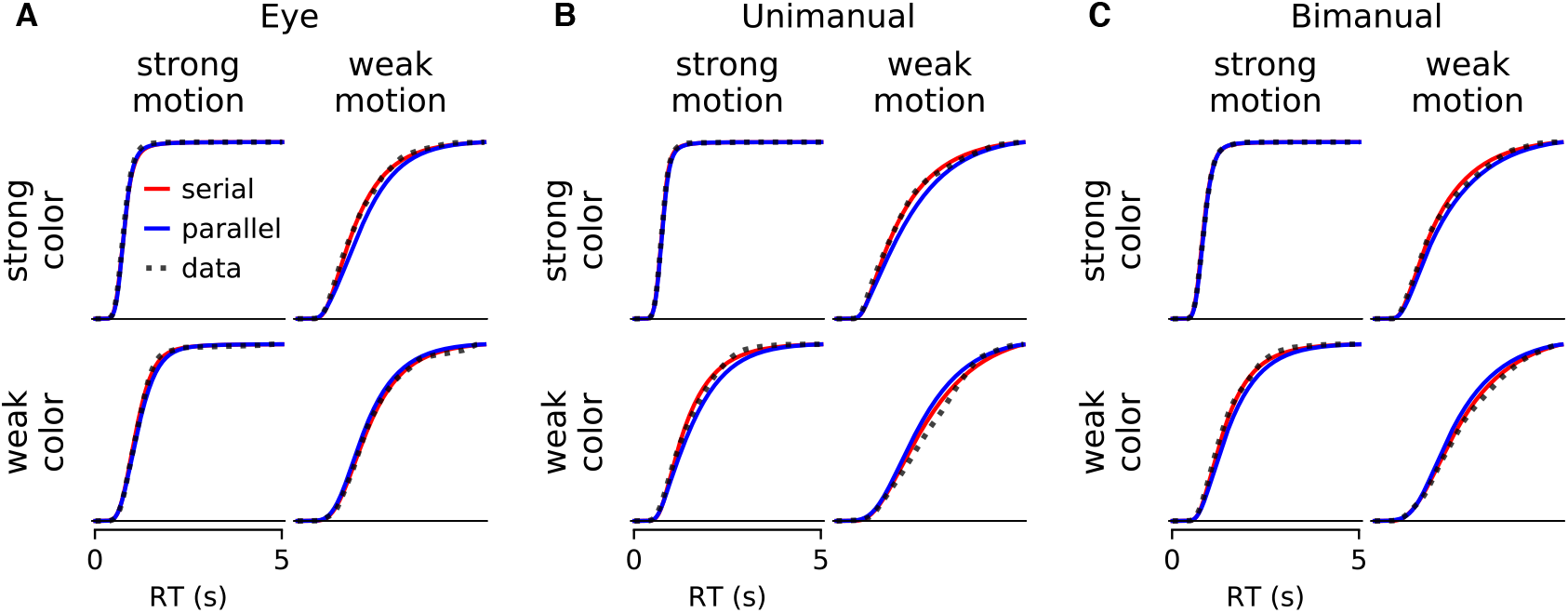
Comparison of parallel and serial rules applied to reaction time distributions. The graphs show averages of the fitted distributions (thick colored traces) across participants. **A**. 3 participants who responded with an eye movement to one of four targets. **B**. 8 participants who responded with a hand movement to one of four targets. **C**. The same 8 participants who responded with two hands (the RT is the time of the last movement). The averages are taken at each time bin across participants for each condition, weighted by the number of trials. Only the conditions with the weakest and strongest stimulus strengths are shown. The comparison provides strong support for the serial combination rule (see ***Figure 2–Figure Supplement 3***).

**Figure 2–Figure supplement 3.**
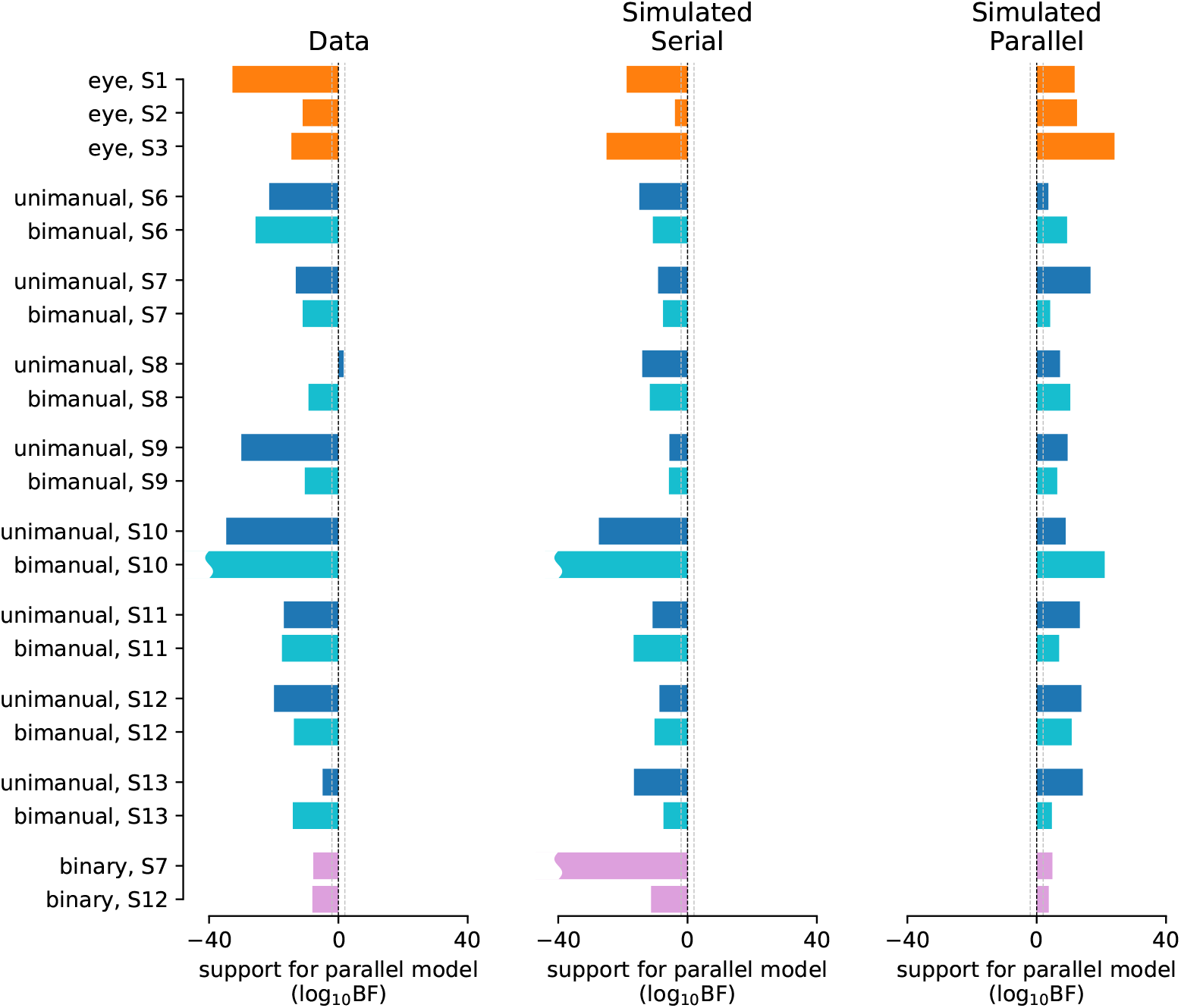
Statistical comparison of parallel and serial rules applied to reaction time distributions. The analysis focuses on the full set of RT distributions, exemplified in ***Figure 2–Figure Supplement 2***. The results are presented in the same format as ***Figure 2–Figure Supplement 1***. Average log_10_BF ± SEM are -17±2, -16±2, 10±1 for the data, simulated serial, and simulated parallel, respectively. For the binary-choice task (pink), the simplified version of the RT model was used and 20 simulations were performed for each participant under the serial and parallel rule, respectively (bars represent the mean across the 20 simulations). For the remaining data, the full RT model was used and only a single serial/parallel simulation was performed for each participant.

**Figure 2–Figure supplement 4.**
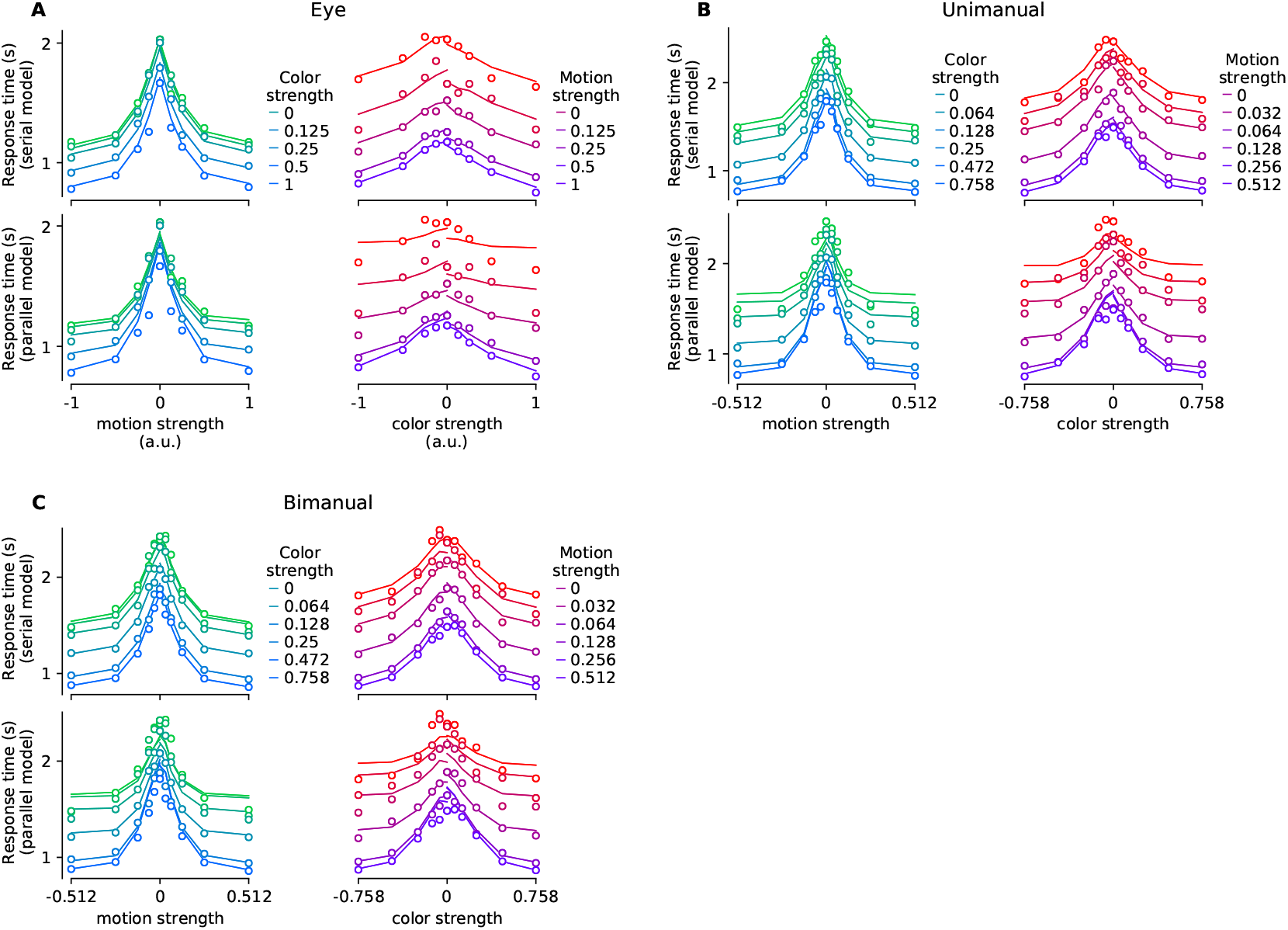
Mean reaction time for parallel and serial rules applied to the reaction time distribution analysis exemplified in ***Figure 2–Figure Supplement 2***. The graphs display the mean RTs and fits in the same format as Fig. 2, with responses reported by eye (**A**), unimanually (**B**), or bimanually (**C**). Mean RTs are computed from the average RT distribution computed as in ***Figure 2–Figure Supplement 2*** for correct choice trials within each condition (or for the zero stimulus strength condition, all trials). Note that these averages across time bins and across participants are used for visualization only; fits were performed for individual participants using the full RT distribution. Here, the fits are derived from the best fitting gamma distributions, described in association with ***Figure 2–Figure Supplement 3***. Open symbols are the data; the traces are line segments connecting the fitted means. In each panel of four graphs, the upper and lower pair of graphs show fits to the serial and parallel models, respectively. Panels display data from the double-decision RT tasks using the three response modalities as indicated.

**Figure 2–Figure supplement 5.**
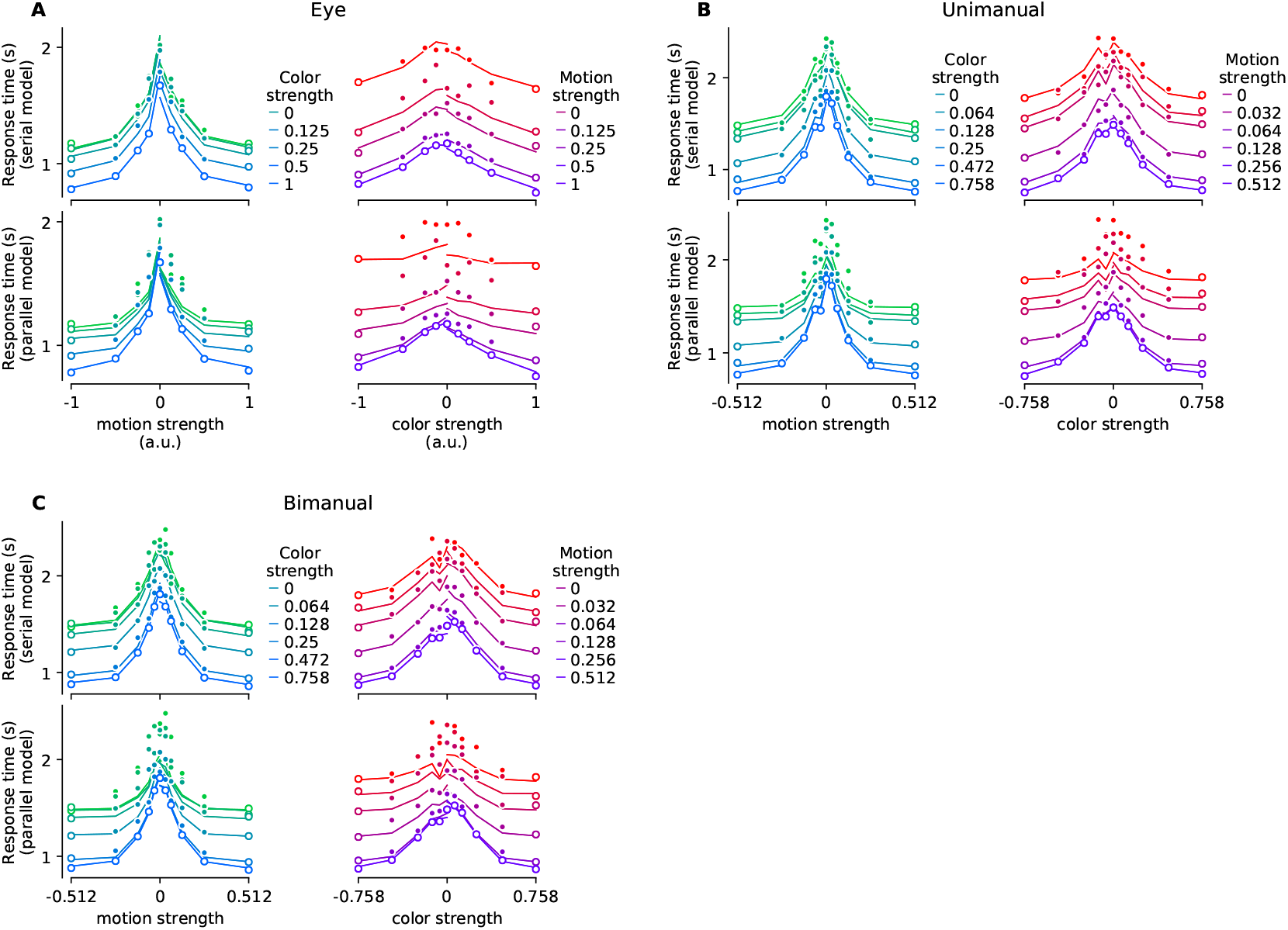
Mean reaction time for parallel and serial rules applied to reaction time distribution analysis with the fit-prediction approach. Format is identical to ***Figure 2–Figure Supplement 4***, except the fits of the marginal 1D distributions were obtained using only the conditions where color or motion strength was at its strongest level. The symbols corresponding to these ‘fitted’ conditions are open. Where the symbols are solid, the data are not fit, but predicted by the serial or parallel logic (traces). Responses were reported by eye (**A**), unimanually (**B**), or bimanually (**C**).

**Figure 4–Figure supplement 1.**
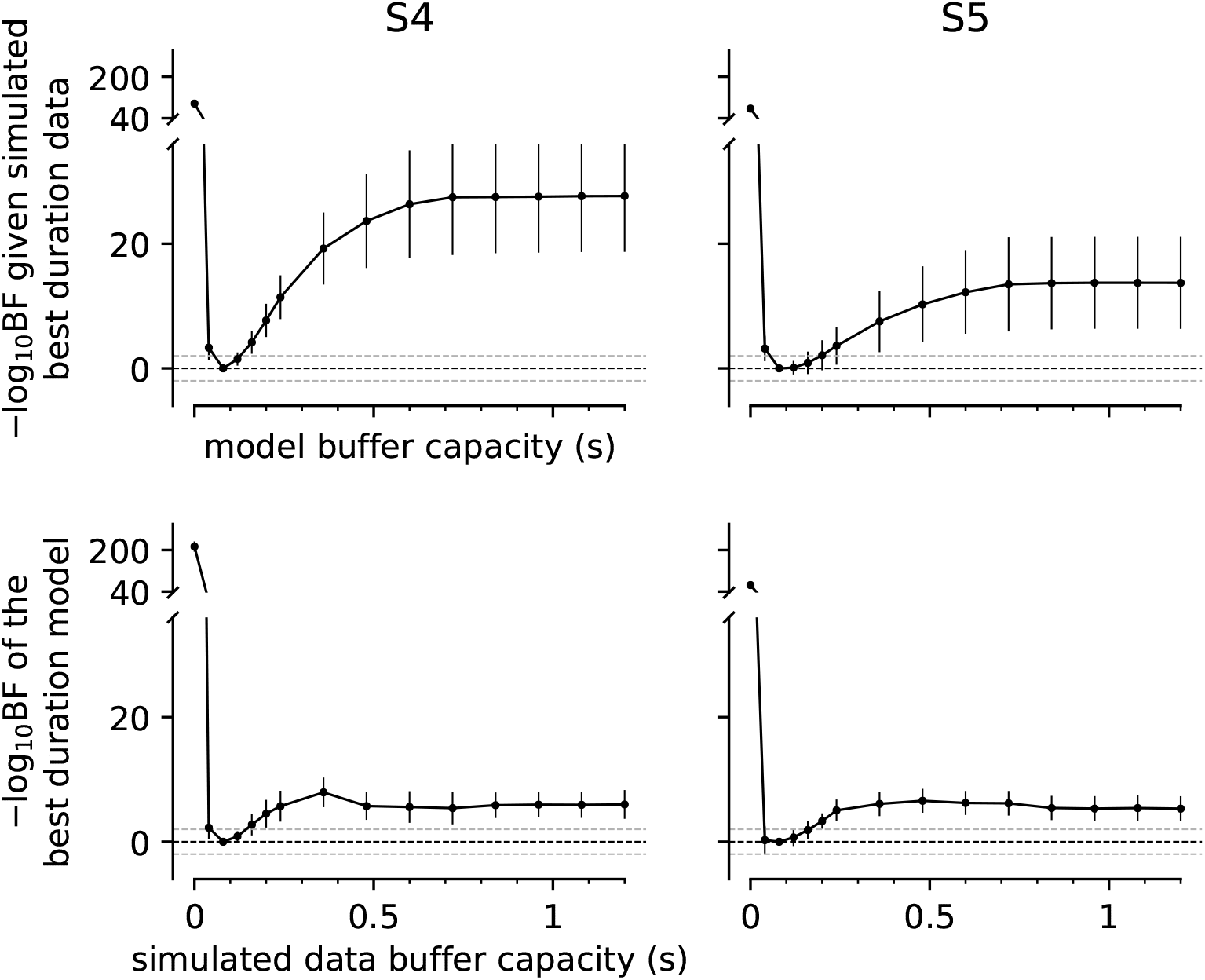
Parameter recovery analysis. The graphs evaluate the sensitivity and specificity of the estimates of buffer capacity (*T*_buf_) shown in Fig. 4. Columns are the two participants. We used the parameters of the best fitting diffusion models to the data in Fig. 4 (solid curves; see Table 5). The analysis in the top row addresses specificity. The simulations use 80 ms, but the model fits used *T*_buf_ fixed to each of the durations shown on the abscissa, computed for a discrete set of buffer capacities (black points). The ordinate shows the difference of each model’s negative log likelihood from that of the 80-ms buffer model (smaller is better). Error bars are standard deviations across 12 simulations. Gray dashed lines show where the Bayes factor = 100 and 1/100 (“decisive” evidence for the best fit model compared to the models above the top line and against the best fit model below the bottom line; ***Kass and Raftery 1995***). The analysis suggests fiducial confidence limits of roughly 80-200 ms. The analysis in the bottom row addresses identifiability. The simulations use *T*_buf_ shown on the abscissa. We then compare two fits, using *T*_buf_ = 80ms or the simulated value. Misidentification is limited to a narrow range similar to the fiducial confidence interval.

**Figure 4–Figure supplement 2.**
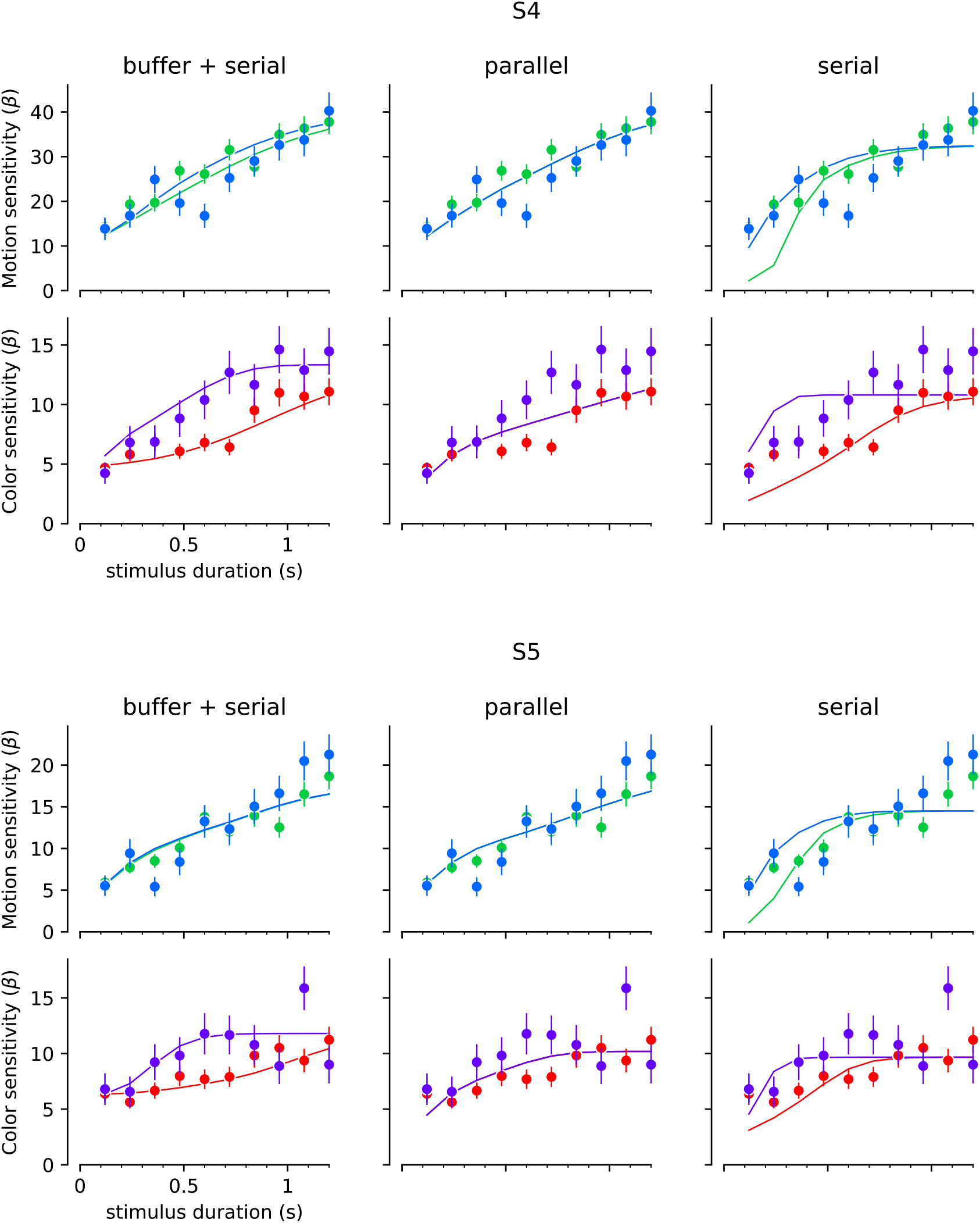
Fits to the choice data with strictly serial and parallel models. The best fitting model to the choice data in the variable duration task implicates a finite buffer, allowing motion or color information to be held for a period before updating the decision. If *T*_buf_ = ∞ or 0, the model is purely parallel or purely serial. The graphs show the best fits of these models for two subjects. The format of the graphs is identical to Fig. 4. *Left column*, reproduction of the fits in Fig. 4. *Middle column*, best fitting parallel model. *Right column*, best fitting serial model. Two participants (rows) performed the color-motion double-decision task with a random dot display that varied in duration between 120 and 1200 ms. *Top*, Motion sensitivity as a function of stimulus duration and color strength. Symbols are the slope of a logistic fit of the proportion of rightward choices as a function of signed motion strength, for each stimulus duration. Data are split by whether the color strength was strong (blue) or weak (green). Error bars are s.e. *Bottom*. Analogous color-sensitivity split by whether the motion strength was strong (purple) or weak (red). Curves are fits to the data from each participant using two bounded drift diffusion models that operate serially after an initial stage of parallel acquisition, here termed the buffer capacity. During the serial phase, one of the dimensions is prioritized until it terminates. The prioritization favored motion for both participants (*p*_motion-1st_ = 0.80 and 0.96, for participants S4 and S5, respectively)

**Figure 5–Figure supplement 1.**
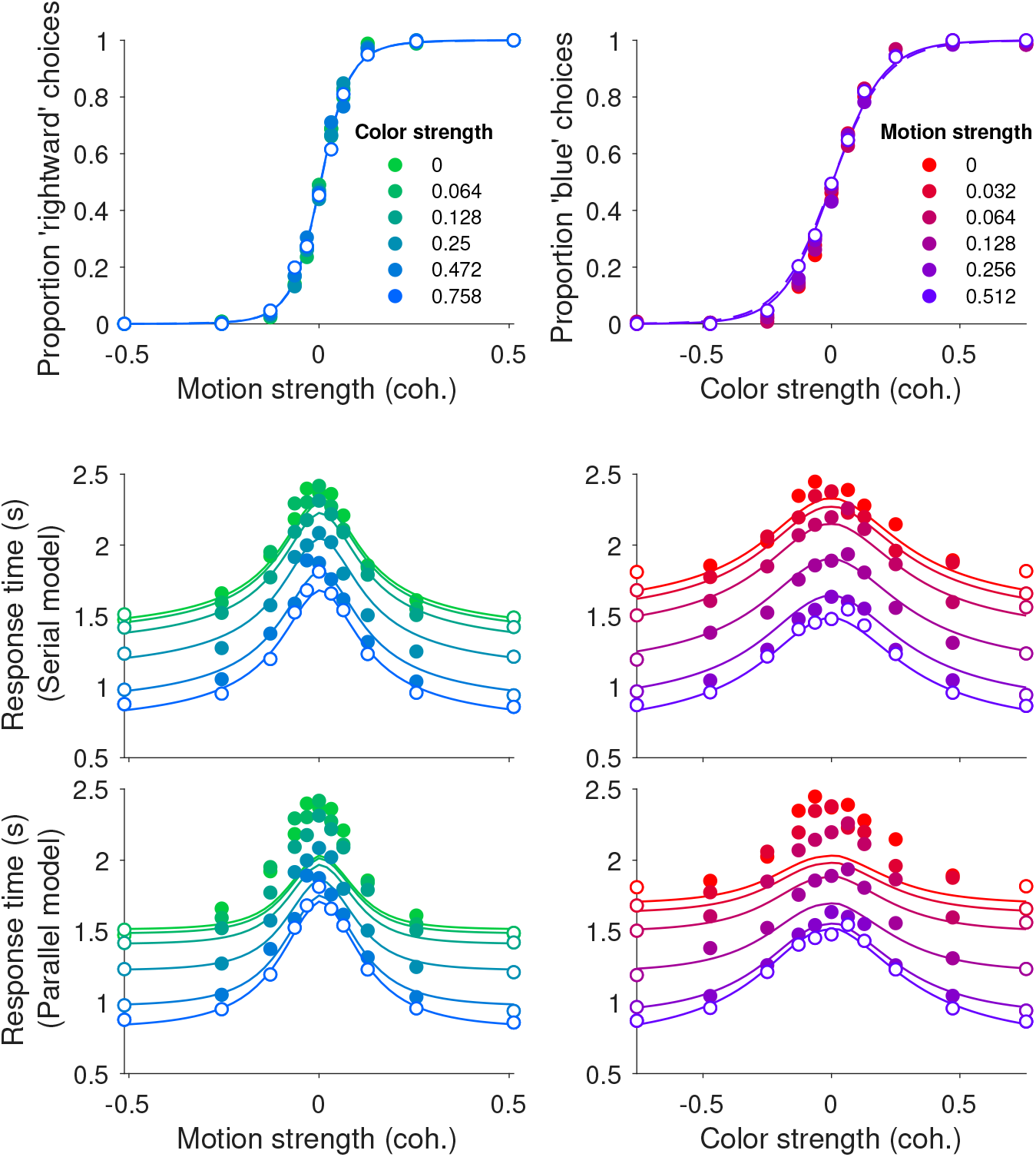
Choice and double-decision reaction time for the bimanual responses in the same format as Fig. 2B. These are the same data shown in Fig. 5 but replacing the predictions from the unimanual fits with the fits to the data from the bimanual task. We use the same fit/prediction strategy as in Fig. 2B. The model comparison summarized in ***Figure 2–Figure Supplement 1***.

**Figure 5–Figure supplement 2.**
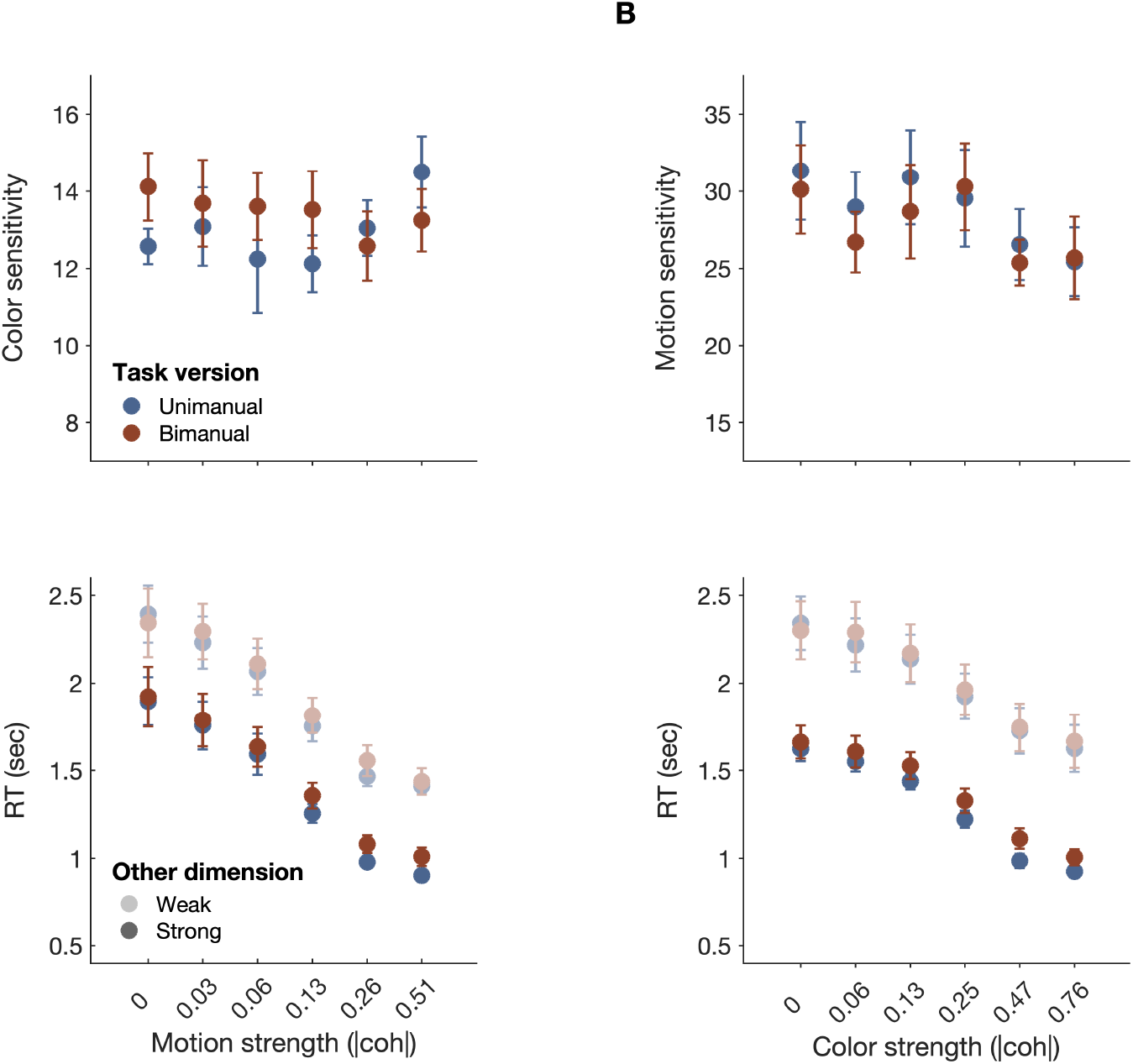
Model-free comparison of performance in the unimanual (blue) vs. bimanual (red) task. **A.** *Top:* Sensitivity of color choices as a function of motion strength (absolute coherence). Sensitivity is the slope of a logistic regression of color choice as a function of signed color coherence, obtained separately for each level of motion strength. *Bottom:* RTs in the uni- vs. bimanual task as a function of absolute motion strength when color was weak (3 lowest strengths; light shading) vs. strong (3 highest strengths;dark shading). For the bimanual task, RTs correspond to the final response of a given trial. **B.** Similar to **A**, but with color strength on the abscissa. *Top:* motion sensitivity. *Bottom:* RTs as a function of absolute color strength when motion was either weak (light shading) or strong (dark shading). No differences in overall choice sensitivity were found between the uni- and bimanual task (repeated-measures ANOVA, motion sensitivity: F_1,7_ = 0.21, p = 0.664;color sensitivity: F_1,7_ = 0.70, p = 0.431). Similarly, overall RTs were similar in the uni- and bimanual task (motion: F_1,7_ = 0.56, p = 0.477;color: F_1,7_ = 0.57, p = .476). Furthermore, the modulation of RTs by the informative and uninformative dimensions, respectively, was not affected by task (uni-/bimanual;all interactions *p* > 0.05). This suggests that overall performance, and modulation of RTs by each decision dimension, were similar in the uni- and bimanual tasks. Data points represent mean ± s.e.m. (N = 8).

**Figure 7–Figure supplement 1.**
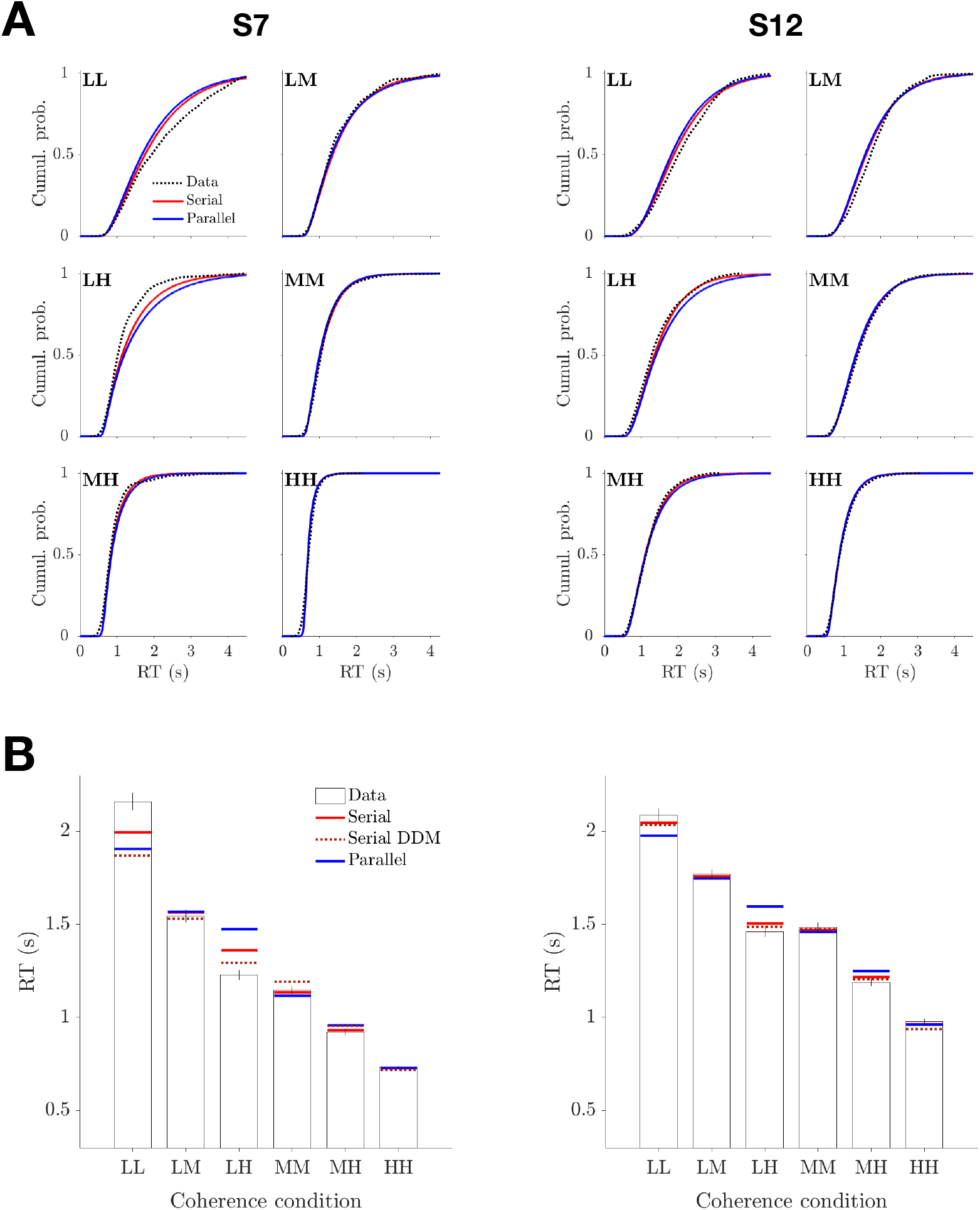
Comparison of parallel and serial rules applied to reaction time distributions in the Same vs. Different task. The analysis is a variant of the one introduced in ***Figure 2–Figure Supplement 2***, applied to RT distributions associated with the six unique combinations of motion strength (correct choices only). The analysis optimizes the parameters of gamma distributions representing three 1D decision times, corresponding to the three unique motion strengths, and one non-decision time to best explain the six observed distributions of RTs. **A**. Best fitting RT distributions for each participant, shown as cumulative probability distributions. Dashed black curves are data. Solid curves are best fitting distributions under serial (red) and parallel (blue) combination rules. **B**. Superposition of the expectations obtained from the fitted distributions (panel A) on the mean RT and DDM fits shown in Fig. 7B).

